# A plant-based biostimulant modulates grapevine susceptibility within a realistic water stress window through priming and phenylpropanoid pathway regulation

**DOI:** 10.64898/2026.02.23.707262

**Authors:** Théo Poucet, Gwennaëlle Chen, Julie Bourg, Anne-marie Busuttil, Chloé E. L. Delmas, Marie-Cécile Dufour

**Affiliations:** French national Research institute for Agriculture, Food and environment, University of Bordeaux, Biology of Fruit and Pathogens laboratory, UMR 1332, F- 33140, Villenave d’Ornon, France; Axioma, 8 Boulevard Voltaire 19100 Brive-La-Gaillarde; INRAE, ISVV, Bordeaux Sciences Agro, SAVE, 33140 Villenave d’Ornon, France

**Keywords:** Abiotic stress, Biostimulants, grapevine, hydraulic conductance, priming

## Abstract

Fluctuating extreme weather events, coupled with rising average temperatures, can severely impact grapevine physiology and yield. While biostimulants have been gaining acceptance as a short-terms tools to enhance grapevine resilience, their adoption is hindered by inconsistent efficacy, partly driven by unpredictable plant stress levels. Over two contrasting seasons, we integrated physiological, transcriptomic, and metabolomic analyses to investigate how a plant-based biostimulant modulates the sensibility of *Vitis vinifera* under varying intensities of heat, drought, and their combination. This panel of water status, ranging from -0.02 to -1.6 MPa, revealed that the physiological response induced by the biostimulant treatment alleviates water stress within a field-relevant hydraulic window located between -0.4 and -1.2 MPa. Moreover, moderate but constitutive reduction of growth parameters in biostimulant plants, suggests a trade-off between vegetative development and abiotic stress responses. Accordingly, gene expression analysis revealed an interaction between water availability and the plant response to the biostimulant, which suggest an activation of priming mechanisms. Metabolic profiling supported these findings, highlighting the central role of phenylpropanoid pathway modulation, together with adjustments in ROS dynamics and stress-related hormone responses, particularly abscisic acid. Overall, this work emphasizes the need for integrating detailed plant water status and leaf gas exchange to accurately evaluate biostimulant performances under abiotic stress.

## Introduction

Climate change is profoundly reshaping agricultural systems, exposing crops to increasingly frequent and severe episodes of drought, heat waves, and their combination. While each individual stress may have only a minor effect on plant growth and survival, their combined application results in a increasing detrimental impact on the plants^1^. Grapevine, one of the most economically valuable crops in the world^2^ is vulnerable to fluctuations in water availability and rising temperatures, which directly affect photosynthetic efficiency, growth dynamics, berry composition, and ultimately yield stability and wine quality^3^. Ensuring grapevine resilience under these conditions is therefore a critical challenge, driving the search for innovative and sustainable strategies that can complement conventional vineyard management.

In this context, plant biostimulants have emerged as promising tools to enhance crop tolerance to abiotic stresses in an environmentally sustainable manner. By stimulating plant growth, physiological and metabolic processes rather than directly supplying nutrients, biostimulants have been shown to improve nutrient use efficiency, modulate hormonal balance, and reinforce tolerance to abiotic stresses such as drought and heat stress^4^. Commonly, 6 subcategories of non-microbial plant biostimulants are distinguished: chitosan, humic and fulvic acids, animal and vegetal protein hydrolysates, phosphites, seaweed extracts, and silicon^5^. More recently, an additional group of plant biostimulants, the plant-based extracts, has received significant attention. They can be derived from a wide range of plant species^6^ and are relatively easy to prepare. Even if this is dependent of the extraction process, these extracts are rich in structurally diverse bioactive molecules, including phenolic acids, flavonoids, terpenes, alkaloids, and other secondary metabolites. These compounds have been reported to regulate antioxidant activity, modulate signaling pathways, and contribute to enhanced plant resilience under stressful conditions. Several studies in model plants and crops such as tomato, maize, and grapevine among others, have demonstrated their potential in alleviating the detrimental effects of salinity, water deficit or high temperature by improving physiological performance and reducing oxidative damage^7,8^. Accordingly, a comprehensive meta-analysis of 180 studies worldwide on biostimulants has classified plant extracts as the category with the best performance in terms of crop yield response^9^.

Despite this potential, plant responses to biostimulants, are often highly variable and context-dependent. This variability reflects several sources of complexity. Their chemical composition is intrinsically heterogeneous, depending on the botanical species used, its phenological stage, the type of tissue collected and the extraction process^8^. Application practices further add to this variability, since plant phenology, dosage, frequency, and delivery mode (foliar versus root) can all lead to contrasting plant responses^9^, especially when extracts are applied either as a priming strategy before stress, or as a mitigating treatment during stress. Plant responsiveness itself represents another layer of variability: effects are rarely uniform across species^10^ and within a single crop, differences among cultivars or rootstocks in basal stress tolerance and physiological plasticity strongly influence outcomes^11,12^. Beyond these biological and technical factors, the environmental and climatic context exerts a dominant effect^11,13^, yet our understanding of the precise windows of efficacy of the biostimulant remains limited. Even when multi-year data are available, variability is often reduced to a description of the conditions under which plants developed, without fully characterizing the intensity, duration, or dynamics of the stress to which they were exposed. As a result, integration of stress characterization into experimental designs is still rare, hindering our ability to link biostimulant responses to specific stress scenarios and to decipher the mechanisms underlying their action.

In this context, the present study aimed to investigate the effects of a complex biostimulant, composed of multiple plant-derived fractions, on grapevine responses to drought, heat, and their combination. To capture the inherent variability of field conditions, experiments were conducted under semi-controlled conditions over two successive years that differed markedly in their weather conditions, particularly in terms of temperature, which resulted in contrasting intensities of water deficit. Nevertheless, through systematic monitoring of plant water status in the different combinations of stress, we characterized a continuum of drought intensity across experiments, providing a robust framework to evaluate treatment effects. This approach not only enabled us to demonstrate that the biostimulant reduces the susceptibility to water, but also delineate a window of efficacy under semi-realistic conditions. Furthermore, transcriptomic and metabolomic characterizations revealed a priming effect of the treatment, notably through the modulation of the phenylpropanoid pathway in a tissue-specific manner, leading to divergent responses in roots and leaves. These molecular and metabolic adjustments suggest a trade-off between organ growth and stress preparation, ultimately contributing to improved tolerance to water deficit. Together, these findings provide new insights into the mechanisms of action of a biostimulant in perennial crops and highlight the importance of integrating stress characterization into the evaluation of biostimulant efficacy.

## Results

Over two independent experiments performed during 2021 and 2022 seasons, a solid form of a plant-based biostimulant was applied to the root system of grapevine plants at leaf senescence in late fall, while the liquid form of the product was sprayed on shoot three times, when leaves are fully expanded, at floral bud emergence, and at the fruit set stages. Then, control and treated plants were exposed to control conditions (no stress, NS), Heat (H), Drought (D) and combined stress (H:D) under semi-controlled conditions (Fig.1).

**Figure 1:**
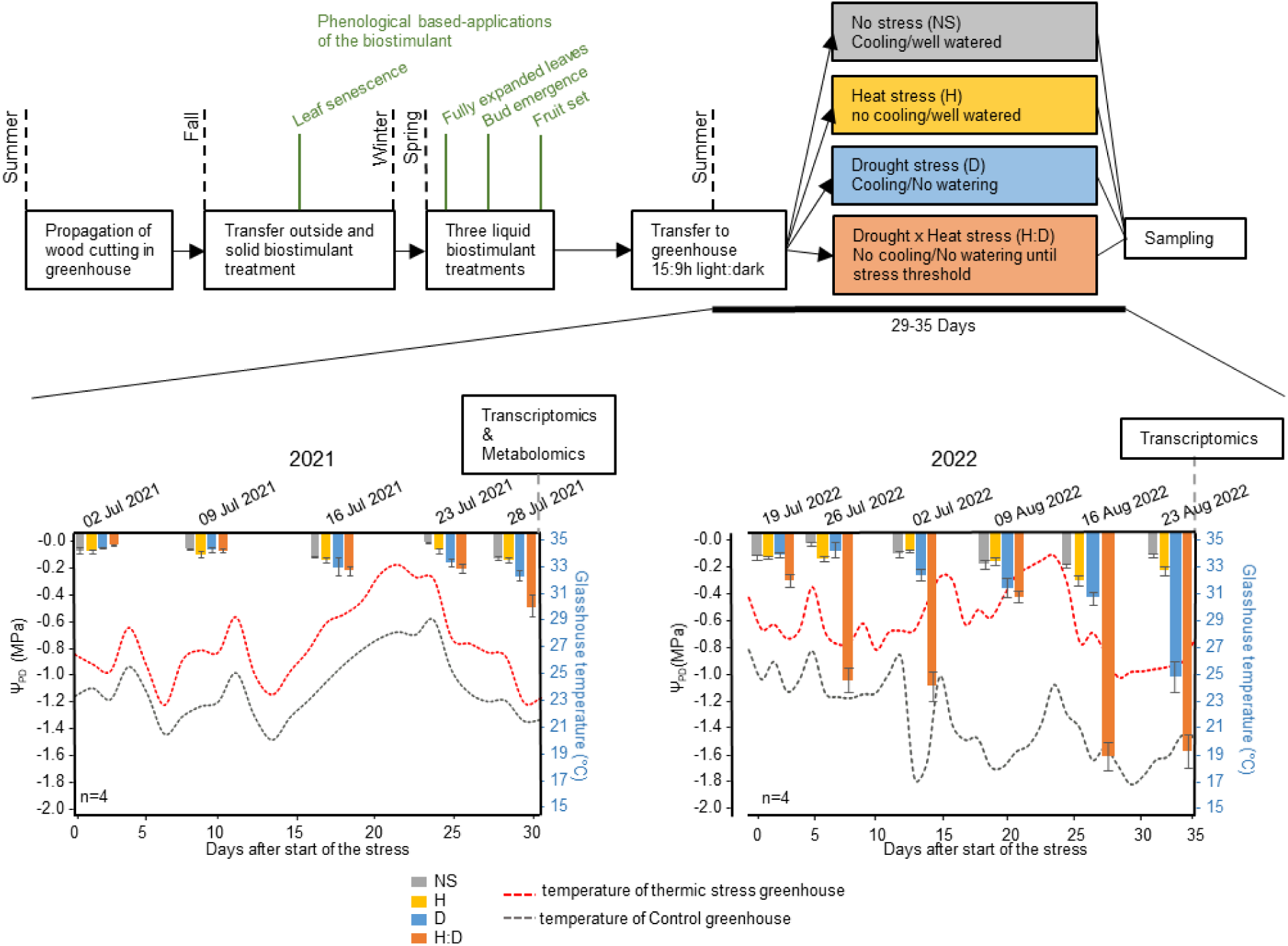
Procedures and stress monitoring for grapevine experiments conducted in 2021 and 2022. Grapevines plants were propagated and kept outdoor. Solid biostimulant treatments were applied before leaf senescence at fall. After reappearance of leaves during spring, three liquid treatments of roots were applied, when leaves are fully expanded, at floral bud emergence, and at the fruit set stages. During summer, plants were transferred to the greenhouse (15:9 h light: dark), under semi-controlled conditions for 29-35 days: No Stress condition (Grey, cooling system/Well watered), Heat stress (yellow, no cooling system/Well watered), Drought stress (blue, cooling/no watering) and Drought combined with Heat stress (orange, no cooling/no watering). Plant stress level was monitored through Ψ_PD_ (n=4 ±se). Stress threshold was established between -0.5 and -0.6 Mpa. Green and red dashed lines indicate averaged temperatures in the two different greenhouses with or without temperature cooling system leading to no thermic stress condition (NS and D conditions) and thermic stress condition (H and H:D conditions), respectively.

### Treated plants show moderate shoot and root growth reduction under mild water deficit but preserve root biomass under severe stress

Without considering treatment effects, under NS conditions and H stress, predawn water potential (Ψ_PD_) was maintained at a constant level (-0.152 MPa ±0.01) for both years, whereas it decreased over the time under drought conditions. Especially under combined stress, that decrease was much faster in 2022 (up to -1.6 MPa ±-0.18) compared with 2021 (up to -0.53 MPa ±-0.09). Reported temperature in cooled (NS and D conditions) vs no cooled (H and H:D conditions) greenhouses showed longer and higher differences in 2022 compared with 2021 year. Overall, such temperature differences, together with lower Ψ_PD_, indicates that plants from drought modalities in 2022 suffered a more intense drought stress compared with 2021 respective plants (Fig.1).

Physiological parameters were evaluated at the end of stress application. According to ANOVAs, biostimulant treatments significantly impacted shoot length, number of internodes and root weight in both years (Fig.2a). Globally, number of internodes and shoot length were slightly reduced under biostimulant treatment compared with control conditions and independently of drought or heat stress effects. Total leaf area (evaluated in 2022), and chlorophyll contents were not significantly affected by the biostimulant treatments. Leaf area was rather stable under NS, D and H stress conditions, but a significant drop occurred under H:D condition. On the other hand, no change of chlorophyll contents was reported in 2021, whereas heated modalities of 2022 (H and H:D) showed higher accumulation. Importantly, we observed a slight but constitutive significant decrease of root biomass in treated plant in 2021, independently of stress modalities. This is illustrated with highly significant Treatment (T) effect according to ANOVA tests. By contrast, root biomass accumulation showed a differential behaviour in 2022, with the appearance of interaction effects between Treatment and Stress variables (TxS). Indeed, in 2022, root biomass was significantly reduced in treated plants under NS condition but remain then stable and similar as control plants under single H and D stress. Nevertheless, the combined stress H:D of 2022, previously identified as the most intense water stress conditions of the experiment, leaded to significant and differential drops of root biomass of 63 % for control plants and only 46 % for treated plants, when compared with other stress conditions (Fig.2a).

**Figure 2:**
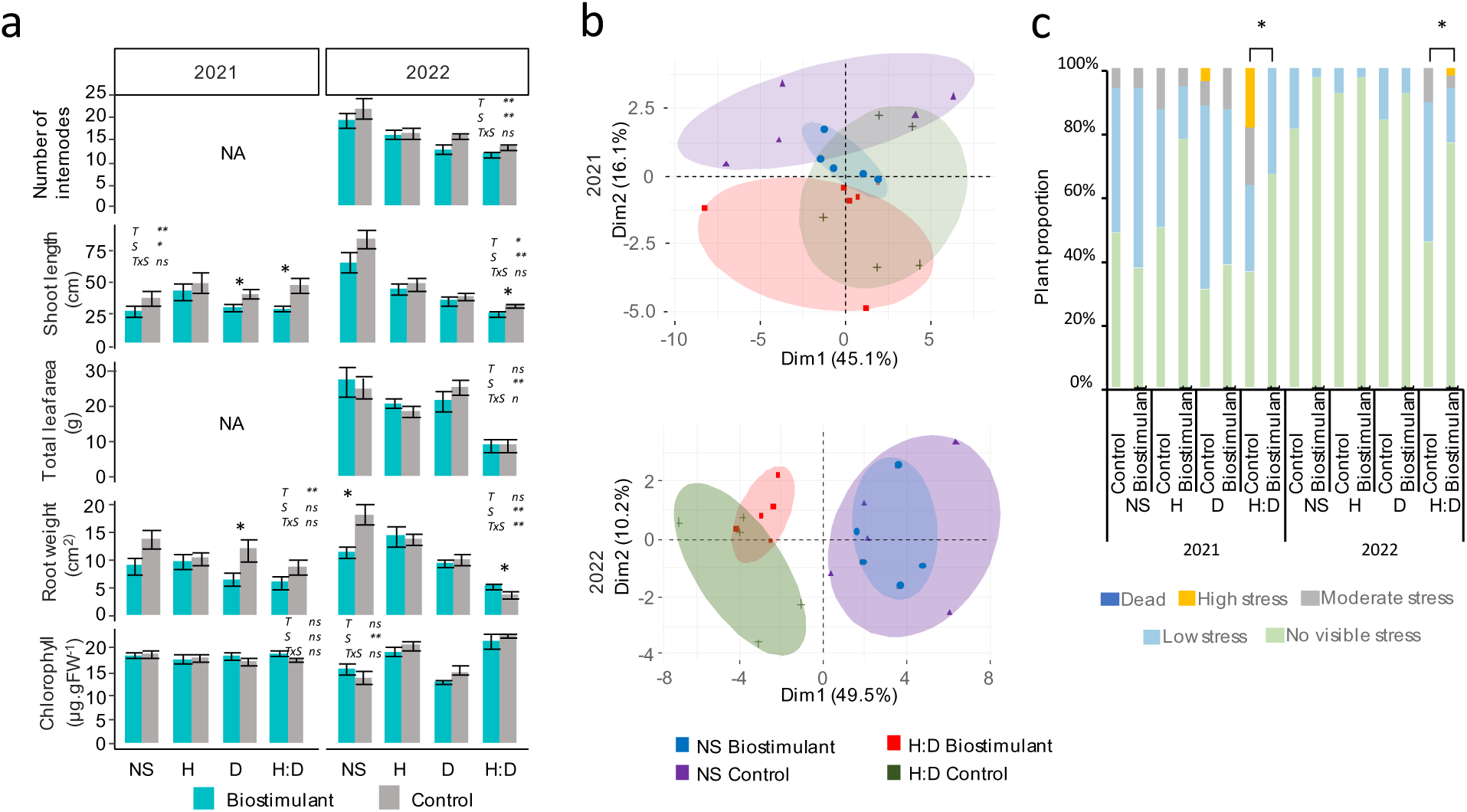
Morphological parameters of Biostimulant (blue) and Control (grey) plants under Heat (H), Drought (D) and combined stress (H:D) during two consecutive years. a) Number of internodes, shoot length, total leaf area, root weight and leaf chlorophyll. Value represent mean ± se (n=10). Significant differences are shown according to two-way ANOVA (**: p<0.01; *: p<0.05; .: p<0.1) where T indicates biostimulant treatments effect (2 modalities: Biostimulant and Control), S stress effect (4 modalites: NS, H, D, HD) and TxS, interaction effect. b) Principal component analysis (PCA) plots of root architectural parameters of Biostimulant and Control plants under no stress and combined stress conditions (H:D) in 2021 and 2022. PCA was performed from the correlation matrix generated with a total of 29 variables (n=5). Plots on the right show the most contributing parameters to the first and second dimensions of PCAs obtained each year. c) Distribution of visual stress groups under control, thermic, drought and combined stress conditions over two years, after 30-35 days of stress. Group Percentage represent At least 10 observations per modalities. Treatment effects on group repartitions were tested with Fisher’s test with 5% threshold. Significant difference between treatments is indicated with an asterisk (pValue<0.05). Illustrative pictures for different stress levels are shown in Figure S2.

Picture analysis of cleaned roots, generated 29 descriptive features of root architecture, used for performing PCAs. Most differences in root architectures were observed by comparing NS with H:D stress, as shown in Figure 2b. Mild discrimination of different modalities occurred in 2021 through PC2 (16.1 %), whereas clearer separation driven by PC1 (49.2 %) occurred in 2022. In the latter, treated roots were architecturally identical under NS control, but were slightly different from control conditions under combined stress conditions, despite a small overlap of the conditions. This difference is partially driven by PC2 (10.2 %), for which the most contributing variables are “Center of mass” (17.8 %) and “Length distribution” (16.8 %; Fig.S1). As an integrator of plant physiology, visible impact of treatments and stresses was evaluated on the plants (Fig. S2 and Fig2c). We found a significant increase in the number of plants in the biostimulant modality without visible stress in H:D conditions compared to the control, in 2021 and in 2022, as well as a reduction or even an absence of plants presenting moderate or high stress.

### A panel/Broad range of water stress levels reveals that the biostimulant induces lower susceptibility to water stress

To better understand if softening water stress pressure takes part to the biostimulant mechanisms, plant water status was monitored with leaf stomatal conductance (*g_s_)*, Ψ_PD_, and midday water potential (Ψ_l_) measurements, over the duration of imposed conditions (Fig.3). *g_s_* showed two distinct patterns over the two years (Fig.3a). First, for both years and according to ANOVAs, treatment and time had no significant impact on unstressed plants (NS) and heated plants. However, *g_s_* remained consistently high for H plants. Secondly, treatment effects on *g_s_* were detected under H:D plants in 2021 and as a Time-Treatment interaction (TxD) under D stress in 2022. In both cases, stomatal aperture of treated plant was maintained opened longer than the control, with significantly higher *g_s_* at 15 and 26 days after start of the stress in 2021 and 2022 respectively (Fig.3a)

**Figure 3:**
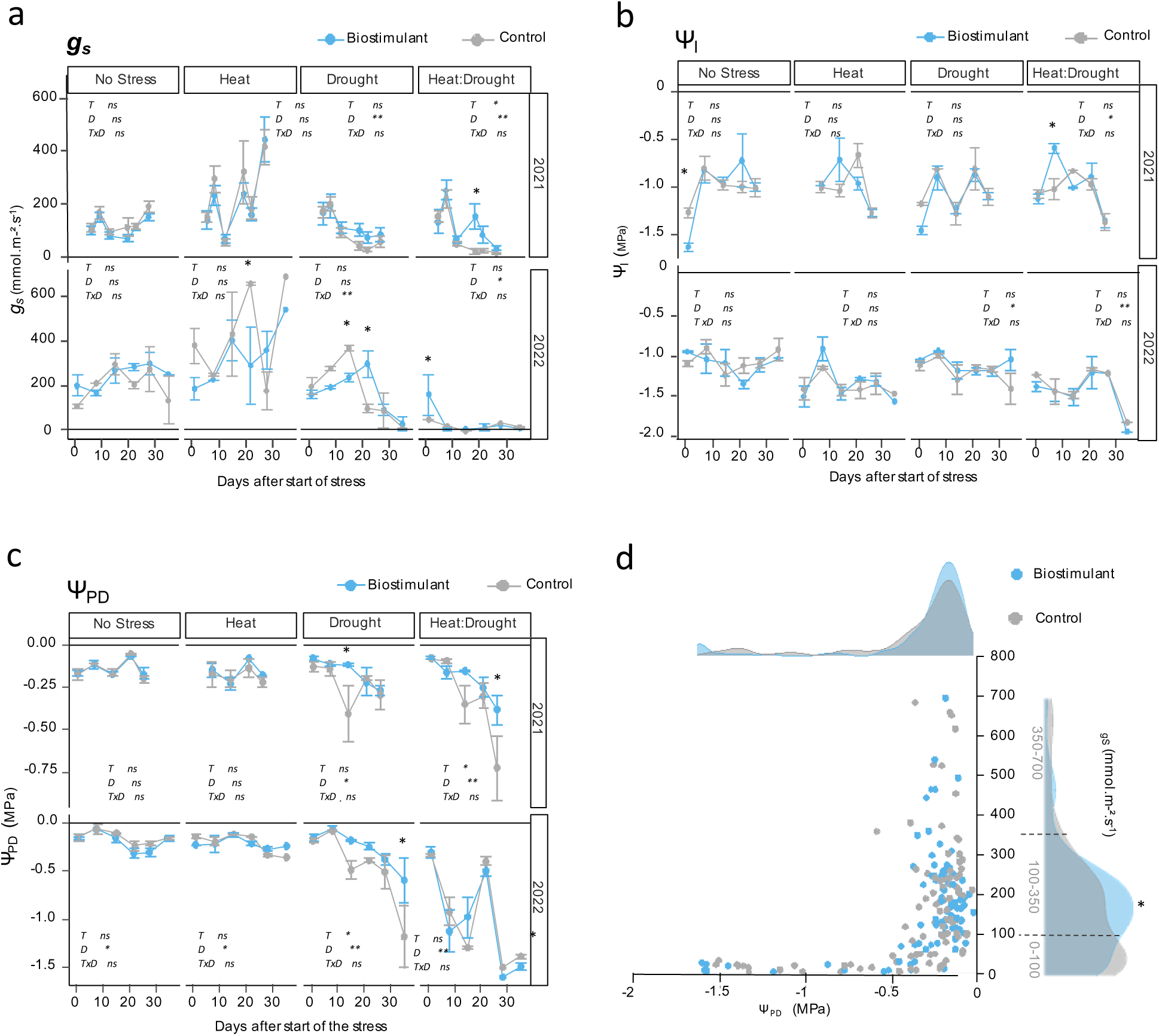
Leaf stomatal conductance and water status of biostimulant treated and control plants under Heat:Drought stress and visual stress scores during two consecutive years. Time-course evolution of stomatal conductance a) g_s_, b) Midday leaf water potential (Ψ_l_) and c) Predawn leaf water potential (Ψ_PD_) of control (grey) and biostimulant (blue) treatment throughout stress applications. Value represent mean ± se (n=3-10). Significant differences are shown according to two-way ANOVA (**:p<0.01; *:p<0.05) where T indicates treatment effect, D days effect and TxD, interaction effect. d) Relationship between Ψ_PD_ and g_s_ under control (grey, n=96) and biostimulant (blue, n=95) treatment. Density plots show the repartition of plant Ψ_PD_ and g_s_ on x and y-axis, using all the measurements performed during the stress application in both 2021 and 2022 seasons pooling together the 4 stress modalities (NS, H, D and H:D). Treatment effects on group repartitions (ranges are indicated in grey of y-axis) were tested with Fisher’s test with 5% threshold. Significant difference between treatments is indicated with an asterisk (pValue<0.05).

Although not significant, it is worth noticing that in 2021, under D condition, the *g_s_* response of treated plants tended to shift (p_T_=0.77; p_D_=0.05; p_TxD_=0.21)) and raised above that of the controls from 15 to 26 days after start of the stress. This trend aligns with a brief but higher significant Ψ_PD_ of treated compared with control plants, observed at 15 days after start of the stress. Finally, under the most severe combined heat and drought stress applied in 2022, we observed that the first time point still showed a significantly higher *g_s_* in treated plants, before to eventually stabilize at similar low levels in both treated and control plants.

For Ψ_l_ parameter and according to ANOVAs, control and treated plants followed quiet similar pattern during the different stress applications (Fig.3b). However, significant impact of the biostimulant treatment on Ψ_PD_ were detected under combined stress H:D in 2021 and D stress of 2022 (Fig.3c). For these conditions, treated plants exhibited slower decrease of Ψ_PD_ over the time when compared with control conditions, leading to significantly higher potentials at the end of the kinetics. It is worth noticing that distinguished potential between control and treated plants start to be visible when the potential of control plants decreases to around -0.4 MPa. Similarly, the lower Ψ_PD_ of control plants, significantly compensated by the treatment was -1.25 MPa (±1.17), observed at the end of D stress of 2022. Importantly, stomatal regulation was not driving differences observed in *g_s_*, since the relationship between *g_s_* and Ψ_PD_ showed no differential behaviour between control and treated plants (Fig.3d). Moreover, density plots show a significant increase of biostimulated plants with *g_s_* ranging from 100-350 mmol.m-².s^-1^, which correlates with an increased proportion of biostimulated plants with modest but higher Ψ_PD_ (from -0.40 to 0 MPa).

### Integration of a continuum of stomatal conductance responses for gene expression analysis reveals an interaction between water availability and biostimulant effects

To understand how treated plants physiologically respond to combined thermic and drought stress, the expression profiles of 89 defence and stress-related genes were determined in leaves and roots in the two years. Genes were arranged into 8 categories, namely, signalling, cell wall synthesis, hormones, homeostasis, redox metabolism, growth and primary metabolism, secondary metabolism, and defence proteins; giving an overview of the main known regulated pathways of grapevine when exposed to heat and drought (Supplementary table S1). Performing a principal component analysis (PCA) with this dataset shows the first principal component (PC1) accounted for 30.2 % of the total variance and underlines the impact of year of experimentation on gene profiles. Given the strong differences in the stress intensities applied to the plants in the two different years, the impact of the treatment was hardly distinguished when year effect was considered as a single binomial explanatory variable (year 2021 or year 2022, Fig.4a). Actually, when testing the effects of the different modality combinations on the gene expression data with ANOVAs, most of the variance is captured by complex interactions of the year together with other modalities. Instead, transcriptional data from both years has to be integrated and compared through a quantitative variable, reflecting the level of water status at which plants were exposed independently of the year. To do so, the area under the curves (AUC) of stomatal conductance (*g_s_*) parameters was calculated for a total of 16 different AUCs, one for each combination of stress modalities, treatment and year. Among the different measured parameters of plant water status, *g_s_* was considered here as the best indicator of drought stress intensity because i) levels and profiles of Ψ_PD_ and Ψ_l_ is unchanged under NS and H stress modalities in both years, making impossible their differentiation using those parameters. On the other hand, *g_s_* conductance indicates a higher stomatal conductance when compared with all other conditions, revealing a different water status of H stress plants. ii) The link between stomatal conductance and Ψ_PD_ is unchanged with the treatment. Therefore, the estimation of the stress intensity through the AUC*_gs_* will not be biased by a treatment-related differential behaviour of stomata regulation and still depends of the plant water status. Then, PCA using AUC*_gs_* levels for colorations of individuals shows the segregation of plants based on their gradual levels of water status, driven mostly by PC1 (Fig.4b).

**Figure 4:**
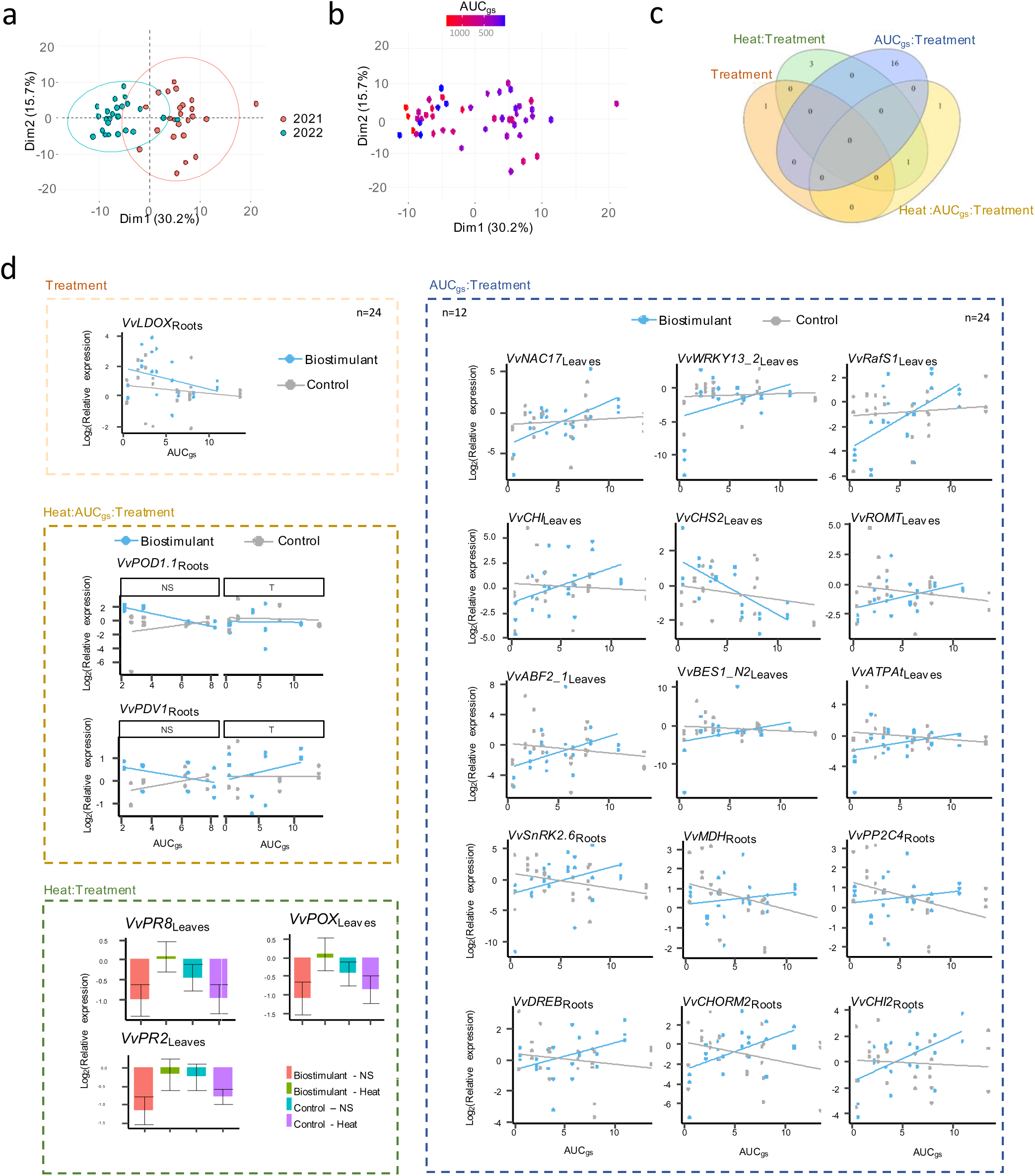
Interaction of biostimulant, heat and AUCg_s_ effect on the expression level of target genes in a two-consecutive year experiment. PCAs (a and b) were performed from the correlation matrix generated with a total of 89 selected gene expressions expressed in root and leaves. Genes were selected according to three-way ANOVA, to evaluate the individual and combined effects of Biostimulant, Heat, and Drought (p<0.05). For Drought effect, AUC (area under the curve) of g_s_ of each modalities was used as a quantitative factor for testing the level of stress that the plants experienced during the trials. Plot (a) shows discrimination by year of experiment and plot (b) shows discrimination by AUC of g_S_ levels. c) Venn diagrams showing the number of genes in leaves or root, significantly affected by the treatment, individually or combined with Heat or Drought factors. d) Relative expression profiles of selected genes (p<0.05) significantly affected by the Treatment factor, individually (Treatment, orange dashed square) or combined with heat (Heat:Treatment, green dashed square), drought stress (Drought:Treatment, blue dashed square) or both (Heat:Drought:Treatment, brown dashed square). For Treatment and Drought:Treatment affected genes, plots show treated and control modalities over AUC of g_S_. For Heat:Treatment affected genes, combined modalities of heat stress and treatment are shown in a barplot. For Heat:Drought:Treatment affected genes, combined modalities of heat stress and treatment are shown over AUC of GSW in a dotplot.

Linear models were built for testing the importance of Heat, AUC*_gs_* and treatment effects on the different genes in leaves and roots as well. In total, 64 genes in leaves and 28 genes in roots were significantly affected by at least one of the previous single factors or combination of them (p<0.05). More precisely, 47 genes in leaves and 17 in roots were exclusively heat and/or AUC*_gs_*-related genes with no effect of the treatment (Fig. S4). In leaves, most of differentially expressed genes were associated with water stress, sometimes with an additional or combined effect of temperature (Fig. S4a). Although fewer genes were differentially expressed in roots overall, temperature, alone or interacting with water deficit, had a proportionally greater impact compared with leaves (Fig.S4b). For instance, only *VvLOX3*, and *VvPGIP* involved in jasmonate biosynthesis and a polygalacturonase inhibiting protein respectively, were exclusively affected by high temperature. Globally, gene expression was negatively correlated with the AUC*_gs_*, meaning that lower water status (lower AUC) led to higher expression levels of target genes. Consistent with this pattern, VvPIP2 and VvTIP2.1, involved with hydraulic adjustments, as well as VvSnRK2.6, VvDREB, VvWRKY30, considered as central genes of dependent and independent abscisic acid (ABA) signalling, constitute a strong molecular signature of adaptation to water deficit. Transcription factors such as *VvABF1* (bZIP family) responded notably to the AUC*_gs_*:Heat interaction, particularly in roots. And other hormone-related genes were upregulated, including those linked to ethylene signalling (e.g., ERFs) or auxin pathways (VvPIN and VvLax2l). Notably, *VvLOX3* in leaves and root appeared specifically linked to the temperature, suggesting a more distinct role for jasmonate signalling in the grapevine heat response.

Concerning the impact of the biostimulant, 21 genes were transcriptionally affected by the treatment or a combination effect of treatment with water stress and/or heat effect (Fig.4d). *VvLDOX*, a gene involved into flavonoid biosynthesis, was the single gene with exclusive treatment effect, as it was constitutively induced in roots in treated plants, when compared with control.

In leaves, *VvPOX*, *VvPR8* and *VvPR2*, all considered as pathogenesis related (PR) genes, showed an interaction effect of the treatment and heat stress. Under control conditions, their relative expression remained low and poorly affected by the temperature whereas the biostimulation triggered significantly higher expression with high temperatures. Fifteen genes were affected by the interaction AUC_gs_:Treatment. Independently of the tissue type, those genes exhibited quiet similar patterns: under control conditions, their expression was rather stable or even increasing with stress intensities, while biostimulation leads to higher expression in NS to mild stress condition, and lower level of expression under hight D stress (low AUC*_gs_*). In this way, the treatment significantly affected major stress signalling genes across different water status, all closely linked to ABA and stomatal regulation, namely *Vv*WRKY13*, Vv*NAC17 and *VvABF2-1* in leaves, and *Vv*SNRK2.6 and *VvPP2C4* in roots. Interestingly, *VvBES1* and *VvDREB* were also influenced, the former being tightly associated with brassinosteroid signaling and growth regulation^14^, and the latter representing a central regulator of ABA-independent abiotic stress responses^15^. *VvRafS1*, known to be involved to raffinose biosynthesis, a trisaccharide largely involved in abiotic stress alleviation as osmoprotector or modulator of redox power among others^16^, was also affected in leaves with AUC*_gs_*:Treatment interaction. Concerning secondary metabolism, regulation and biosynthesis of flavonoid was significantly affected, with *VvCHS2*, *VvCHI* and *VvROMT* in leaves, and *VvCHORM2*, *VvCHI2* and *VvLDOX* in root. Finally, VvPOD1.1, a gene belonging to a peroxidase family involved in redox metabolism and cellular H₂O₂ detoxification^17^, exhibited a distinct expression pattern in roots, in response to varying levels of drought stress, but only under mild temperature conditions. Notably, its expression was reduced in treated plants experiencing D stress (Fig.4d).

### Metabolic characterisation displays tissue-specific adaptation of biostimulated plants under combined H:D stress associated with differential flavonoid accumulations

Transcriptomic data highlighted a treatment-related adaptation of redox and secondary metabolism among others. For this reason, total antioxidant activity assays together with total flavonoid and polyphenol quantifications were conducted on leaf and root tissues across treatments, stress levels, and both years (Fig.5). All three parameters exhibited similar pattern over different tissues and modalities, which highlights the significant contribution of those metabolite families to the antioxidant power of grapevine tissues. However, marked differences were observed between the two years. In 2022, leaves exhibited overall lower levels of reducing power and polyphenols, while roots showed higher values compared to 2021. Notably, ANOVAs revealed significant Treatment effects in leaves of 2021, with a slight decrease of antioxidant power and polyphenol content under biostimulation conditions. No significant changes of leaf Flavonoid contents were detected in leaves. However, in roots, Drought:Treatment interactions were detected in 2021 for FRAP values (near from significance, Pvalue=0.068), and total polyphenol and flavonoid contents (Fig.5). Such interaction effects are explained by a significant increase of all three parameters in treated plant under no stress conditions when compared with the control, while their levels remained unchanged when single or combined stress were applied.

**Figure 5:**
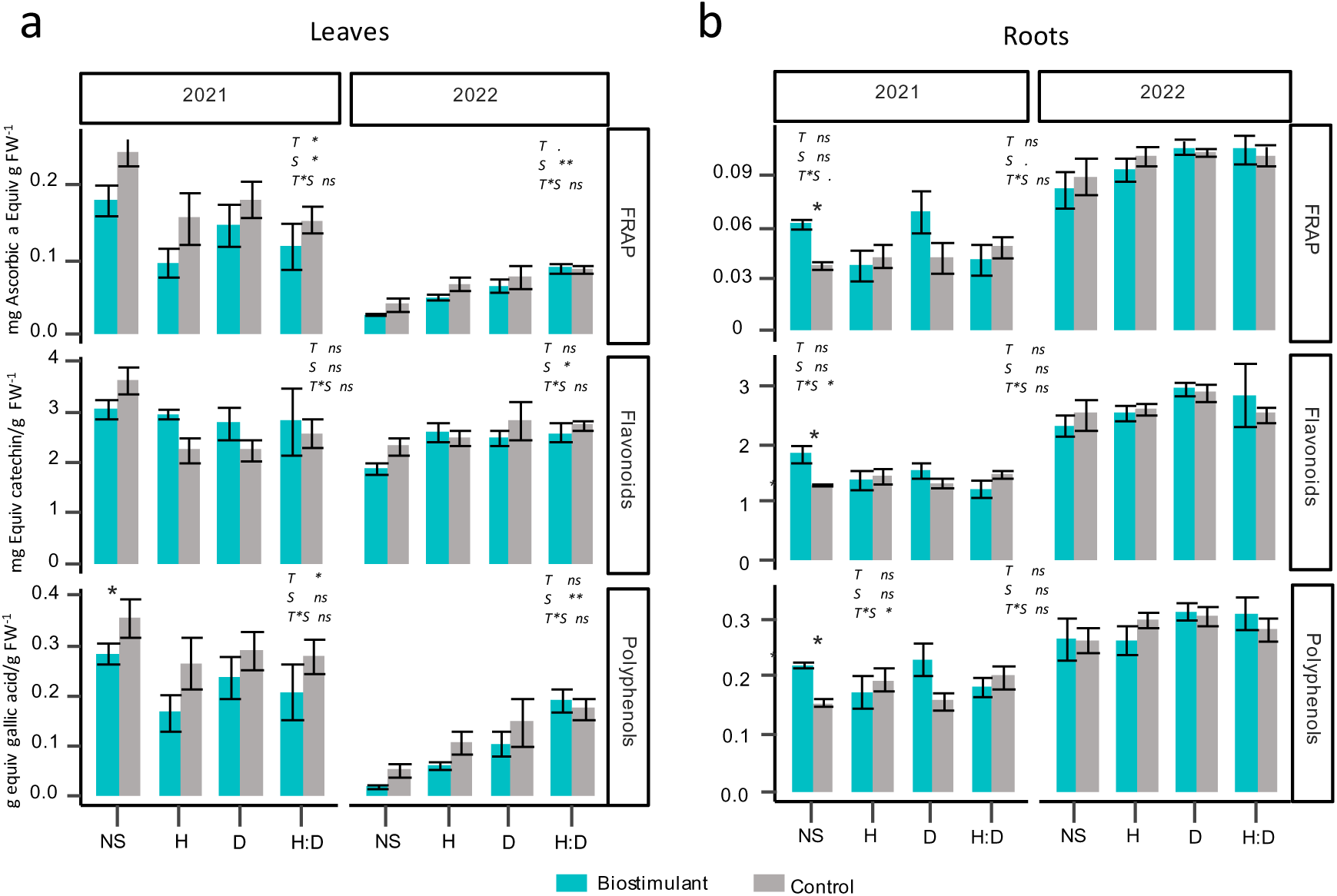
Total polyphenols and Flavonoid content and free radical scavenging activities of treated (blue) and control (grey) plants under Heat (H), Drought (D) and combined stress (H:D) during two consecutive years. a) Leaves and b) roots. Flavonoid, polyphenols contents and FRAP, DPPH scavenging activities. Value represent mean ± se (n=6). Global effect are shown according to two-way ANOVA (**:p<0.01; *:p<0.05; .:p<0.1) where T indicates treatment effect, S stress effect and TxS, interaction effect. Significant differences between control and treated plants are highlighted with asterisks (* p<0.05).

Most contrasting effects between treated and control plants regarding antioxidant metabolites occurred under non-stress conditions. In addition, physiological differences were statistically detectable under mild stress, particularly for water-related parameters. For those reasons, the no-stress and combined heat–drought (H:D) conditions in 2021 were selected for further metabolic characterization. Metabolic profiling was performed through a LC-MS analysis pipeline, providing relative abundance information of 3601 metabolites. In leaves and roots, 700 and 1363 metabolites were respectively detected, as significantly affected by treatment, Heat or Drought, (ANOVA, p<0.05). This indicated a good robustness of the metabolomic dataset with a clear phytochemical diversity shift. Then, unsupervised PCAs and a heatmap analysis (Fig. 6a and 6b) were performed to evaluate overall differences in metabolic profiles of both organs.

**Figure 6:**
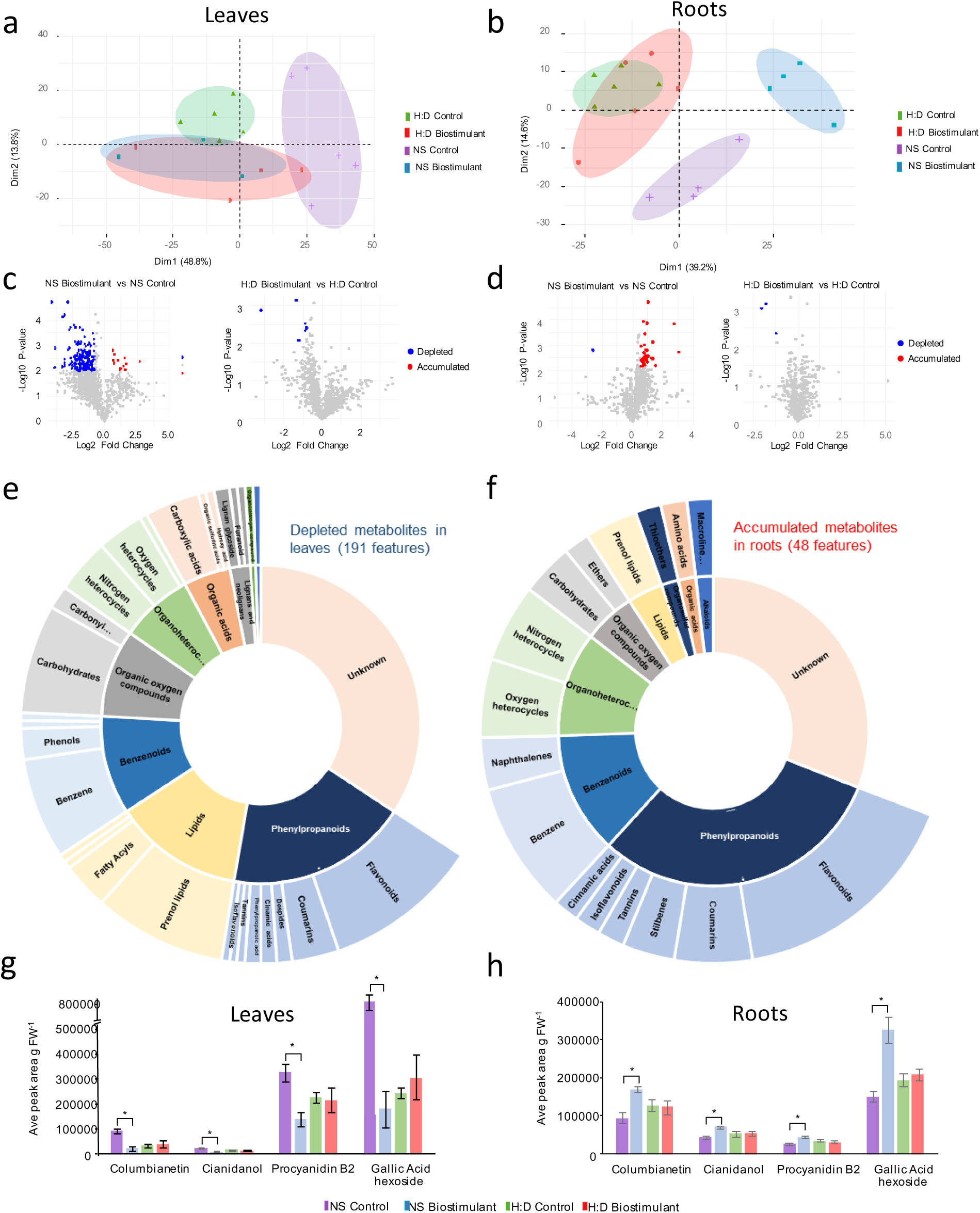
Metabolic profiling analysis of roots and leaves treated with biostimulant under No Stress (NS) and combined Heat:Drought (H:D) conditions in 2021. PCAs were performed from the correlation matrix generated with a total of 1364 metabolites in leaves (a) and 956 metabolites in roots (b), selected according to two-way ANOVA, to evaluate the individual and combined effects of treatment and H:D stress. Volcano plots of metabolic markers that were accumulated (red) or depleted (blue) between biostimulated vs control plants under NS conditions and H:D conditions for Leaves (c) and Roots samples (d). The criteria used for metabolite abundance filtering were α ≤ 0.05, p ≤0.01 and fold change ≥1.5 for upregulated (Red) and ≤0.6 for downregulated (Blue). Sunburst plot of the global metabolome showing the classification (Superclass and Class) of all the putatively annotated metabolites and unknown metabolites depleted under biostimulation in leaves (e) and accumulated under biostimulation in roots (f). Refer to Supplementary Tables 1 for the full names and classification of the metabolites shown in those sunburst plots. Differential quantitative profiles of some of the representative metabolites within main classes of phenylpropanoid pathways impacted by the biostimulant treatment (Tannins, flavonoids and coumarins) and commonly detected in leaves (g) and roots (h). Significant differences between control and treated plants are highlighted with asterisks (* p≤ 0.05).

In leaves, the first principal components (PC1) accounted for 48.8% of the total variance and clearly separated stressed from non-stressed control plants (Fig.6a). In contrast, treated plants displayed highly similar metabolic profiles regardless of stress conditions. Notably, the treated group clustered more closely with the stressed controls along PC1, but were distinguished from them along PC2 (13.8% of total variance). Clustering analysis indicates that such difference between non-stressed controls and biostimulated plants is largely driven by a pronounced reduction in the abundance of numerous metabolites (cluster 1 and 2, from left to right; Fig. S4a). To highlight treatment-related metabolites, pairwise comparison was performed between treated vs control plants from NS and HD modalities. Under NS conditions, biostimulant treatment induced the significant accumulation of 6 metabolites and the depletion of 191 metabolites, while only 6 features were significantly depleted under HD conditions (Fig.6c). Subsequent analysis of the chemical groups of down-accumulated compounds under non-stress (NS) conditions revealed that a substantial proportion belonged to antioxidant-related classes. A large fraction was assigned to flavonoids, including intermediary compounds and derivatives, such as epicatechin (ID:4973), cianidanol (ID: 1968), rutin (ID: 4678), procyanidin B1 (ID: 5482), B2 (ID: 5480) and C1 (ID:4887). In addition, coumarin derivatives such as columbianetin (ID: 4911) was identified, together with trihydroxy benzoisochromen (ID: 2102), an anthraquinone derivative formed from oxidized coumarins. Moreover, tannins such as gallic acid hexoside (ID: 5630) and vulpinic acid (ID: 4241), belonging to furanones, were also detected (Fig.6e).

In roots, the first two principal components (PC1 and PC2), which together explained 53.8 % of the total variance, clearly segregated the treated from control plants under non-stress conditions. Under combined stress, roots showed clear different patterns distinguished from no stress profile. However, biostimulant treatment had no visible impact. Hierarchical clustering showed that treated plant under no stress condition exhibited a marked accumulation of many compounds in the second cluster (from left to right) when compared with all other modalities (Fig. S4B). This cluster was particularly enriched with flavonoid family compounds or derivates (17 %). Overall, 48 features were identified as differentially accumulated, when comparing control and treated plant under no stress conditions (Fig.6d), and 10 of them were classified as flavonoids compounds, derivates or biosynthesis intermediary. Notably, putative flavonoids of particular interest include procyanidin B2 and C1, cianidanol, prunin (ID:3273) or demethyl medicarpin (ID:1353). In addition, other putative polyphenols, such as gallic acid hexoside and the stilbene pinosylvin (ID: 1017), were also differencially accumulated, along with columbianetin and trihydroxy benzoisochromen. Several metabolites of interest for their red-ox buffering role, including cianidanol, procyanidin B1, columbianetin, and gallic acid hexoside, were detected in both leaves and roots (Fig. 6e and f), and illustrate the depletion of antioxidant potential in leaves and activation of redox-related pathways in roots, highlighting organ-specific modulation of the antioxidant metabolism in response to biostimulant treatment under non-stress conditions.

Finally, as major regulators of growth-defence trade-offs and given the previous detection of differentially expressed genes involved in phytohormone biosynthesis pathways, relative abundances of putative salicylic acid, ABA (ID:1410) and jasmonic acid (ID:1012) were specifically examined in the present dataset (Fig. S7). In leaves, the treatment under non-stress conditions triggered a marked increase in salicylic acid levels and a reduction in ABA. On another hand, in roots, treatment had no effect under NS condition but showed lower levels under combined stress conditions, with a significant reduction of putative salicylic acid forms (ID:464). Finally, in both organs, jasmonic acid was no affected by the treatment despite a marked decrease of abundance in roots under stress.

## Discussion

Plant-based extracts are promising solutions for mitigating abiotic stresses such as salinity, heat, and drought^7^. In this study, we investigated the mode of action of a biostimulant derived from plant-based ethanolic extracts of 9 plant species. The biostimulant solid form was applied to grapevine plants as a root treatment at leaf senescence before winter and the liquid form as three foliar sprays at three different phenological stages prior to exposure to individual or combined heat and drought stress (Fig.1). Our results revealed constitutive effects of the treatment on grapevine physiology, independent of the imposed stress conditions. In particular, treated plants displayed a slight reduction in the number of internodes, shoot length, and root biomass (Fig.2). Although these observations may appear as negative growth effects, they did not translate into any reduction in total leaf area, as tested in 2022. Moreover, chlorophyll content and stomatal regulation remained similar between treated and control plants across stress conditions. This suggest that the photosynthetic and transpiration potential of biostimulated plants appeared unaffected.

Importantly, in treated plant the simultaneous maintenance of higher Ψ_PD_ and stomatal conductance (Fig.3d) indicates an improved ability to preserve plant water status and sustain gas exchange under mild water-limited conditions. This suggests that the biostimulant tend to hamper drought susceptibility, likely by supporting root water uptake and/or lower decrease of Ψ_PD_. In the current study, when mild water stress was applied (H:D 2021 and D 2022), we observed the maintenance on average of -0.18 MPa of biostimulated plants when compared to the control. Yet, previous meta-analyses have established a linear relationship between water deficit (expressed as water potential Ψ) and reductions in berry size and yield. For this reason, according to Williams and Phene (2010) and Mirás-Avalos and Intrigliolo (2017), such decrease observed in this study, might on average prevent for loss of around 10% yield^3^.

### Accounting for abiotic stress levels reveals a stress window with optimal biostimulant effects

Variation in the effectiveness of biostimulants is commonly reported, as their impact depends not only on crop species and genotype but also on environmental conditions^9,20^. Over two successive years, monitoring leaf stomatal conductance and hydraulic parameters (*g_s_*, Ψ_PD_ and Ψ_l_, Fig.3) enabled to characterize the plant’s water status precisely across experiments, and to consider a range of stress intensities for defining the window within which the biostimulant was effective. This window first emerged under moderate drought in 2021, when the effects of treatment on *g_s_* were close to significance. Then, *g_s_* and Ψ_PD_ remain significantly higher under both H:D in 2021 and D in 2022. However, when the intensity of stress further increased, as in the severe H:D conditions in 2022, the beneficial effects of the treatment was not detectable, suggesting that the product is effective only within a restricted range of moderate water limitation. It is worth noticing that, although the *g_s_* values were similarly low at midday at the end of the stress period, the significantly higher Ψ_PD_ observed in treated plants suggest that they could have maintained a better water status for a few more days if the stress had continued.

Numerous studies reported a reduced susceptibility of grapevine when treated with biostimulants^21^. Daler and Kaya (2024) showed that the effects of α-lipoic acid were mainly detectable under drought and varied strongly between rootstocks, highlighting the combined influence of stress intensity and genotype on biostimulant efficiency. Whey protein hydrolysate was also showed to efficiently alleviate combined heat and water stress, by preventing dramatic decrease of Ψ_PD_ to an average of -1.75 MPa experienced under control conditions and maintained to -1.23 Mpa when treated^22^. More precisely, Frioni *et al* (2021) reported that a seaweed extract efficiently maintained higher leaf transpiration and water use efficiency when Ψ_l_ values were comprised between -0.6 and -0.8 MPa, while no effect of the treatment could be detected under more severe water deficit. However, such studies were conducted in climatic controlled chambers and compared plant responses to the same progressive and intensifying stress. Alternatively, some grapevine studies include multi-years or environment dimension to their design, and year-to-year variability was acknowledged but rarely linked to plant water status^24,25^. Evidence from tomato further reinforces this view, as multi-year open-field trials consistently reported that the effects of protein hydrolysates or other plant-based biostimulant were strongly modulated by the growing season and irrigation regime^11,13^. In these studies, biostimulants either enhanced productivity or improved fruit metabolic profiles depending on the prevailing weather conditions, while in other years no significant effect was detectable. Such findings underline that biostimulant performance cannot be generalized but must be interpreted in the light of the abiotic stress intensity context.

Irrigated vineyards typically function within a safe range of water potentials, and non-irrigated vineyards rarely reached Ψ_PD_ values inferior to −1.5 MPa, which do not lead to cavitation, turgor loss and/or plant death^26^. Since the efficiency of the biostimulant was observed to be roughly in the range of -0.4 to -1.2 MPa, we can consider that the window of effectiveness of the botanical extract falls within the physiologically relevant range of water potentials for grapevine. By contrast, no impact of the biostimulant was detectable on grapevine physiology when plants exhibited lower Ψ_PD_ values, close to the −1.5 MPa threshold, which represented a situation rarely encountered in vineyard condition. This indicates that the biostimulant remains efficient within a restricted range of water deficit, beyond which the physiological limits of the plant override any potential benefits.

### Biostimulant applications cope with a trade-off between growth and physiological functions under stress through priming mechanism

A significant proportion of genes selected for transcriptomic analysis followed similar and consistent pattern, according to plant’s stomatal aperture, in leaves and to a lesser extent in roots (Fig.S4). Typically, those genes, involved in water and heat stress management, are well known (Table S1 for references) and showed an increasing expression when AUC*_gs_* decreased, which is considered as markers of water availability.

When combined with different stresses, biostimulant treatments caused a subset of key genes to appear, which differed from other marker genes: higher expression under high water abundancy but lower levels when water was lacking and AUC*_gs_* was reduced (Fig. 4). In this way, *VvNAC17* and *VvWRK13*, belonging to two families of transcription factors usually considered as priming markers involved with drought adaptation^4,27^, were significantly affected by the biostimulant. They were often observed with biostimulation-related priming in tomato^28^, grapevine with seaweed extract^29,30^ and cotton with γ-poly-glutamic acid enhancing drought tolerance^31^. In line with these observations, several genes typically associated with stomatal closure under drought stress were down-regulated in biostimulated plants. Notably, *VvRafs2*, a raffinose synthase whose protein activity contributes to drought tolerance in maize and *Arabidopsis*^32^ transcriptionally linked with ABA levels^33^, and *VvABF2*, a bZIP transcription factor implicated both in stomatal regulation and in enhancing cellular drought tolerance^34^, showed reduced expression under stress in treated plants. These transcriptional changes are consistent with the lower ABA levels detected in leaves of treated plants (Fig.6d) and may explain the maintenance of stomatal aperture over the stress periods^35^. In roots, no clear differences in ABA levels were detected between treated and control plants (Fig. S7). Nevertheless, key regulators of ABA signalling such as *VvPP2C* and V*vSnRK2* were affected by the treatment. Notably, the up-regulation of *VvSnRK2* under optimal conditions may prime the root by activating downstream transcription factors^4^, while the concomitant induction of *VvDREB* highlights the reinforcement of both ABA-dependent and ABA-independent stress pathways^15^.

When investigating potential priming effect at the metabolic level, non-stressed (NS) plants cannot be considered strictly naïve. Nevertheless, we can hypothesize that they only experienced the treatment phase, and that they can be considered as a proxy of primed but not triggered plants when compared with stressed conditions^27^. Data indicates that leaves developed a new metabolic profile, which was closer to the stressed state, yet still clearly distinct, independently of cultivation conditions (Fig.6a). This shift was partly driven by the depletion of putative phenolic compounds (Fig.5) including powerful antioxidant molecules such as putative gallic acids, coumarins and various putative flavonoids (Fig.6a). This is consistent with both the reduced total flavonoid content measured in leaves and the modulation of flavonoid biosynthesis genes such as *VvCHI* and *VvCHS2* (Fig.5d). A comparable decline in foliar flavonoids has also been observed in grapevines treated with kaolin^36^. Flavonols are known to exert profound effects on leaf development and physiology; for example, the absence of flavonols in the *tt4-1* (*CHS*) mutant of *Arabidopsis* results in smaller leaf area and slower inflorescence growth ^37^. In line with this, we observed a constitutive and slight decrease in certain biomass traits, such as internode number and shoot length, in treated plants, but no reduction in leaf surface area was detected, at least during the 2022 season (Fig.2). Notably, the observed depletion of flavonoids seems contradictory with their reported role in antagonizing ABA-induced stomatal closure in both *Arabidopsis* and tomato^38^, a discrepancy that warrants further investigation in grapevine.

By contrast, in roots, the biostimulant induced a significant transcriptional and metabolic modulation of the phenylpropanoid pathway, leading mainly to the accumulation of probable flavonoids when no stress was applied (Fig.5 and Fig.6b). Such divergence in the root metabolome was only observed under no-stress conditions. This may suggest that treated roots undergo a preparatory adjustment prior to stress, which is later overridden once stress is imposed. Accordingly, root metabolic profiles visualized through PCA (Fig.6b) showed a clear dissociation only under no stress condition driven by putative flavonoid and precursor accumulations among others. Congruent with metabolic data, *VvCHORM2*, and *VvCHI2*, two genes initiating the biosynthesis pathway of flavonoids through the conversion of chorismate to phenyalanine and p-coumaryl CoA to leucoanthocyanidin respectively, showed higher expression with high AUC*_gs_* compared with control plants, and remained similarly lower with the control under low AUC*_gs_*. Similarly, *VvLDOX*, which catalizes the conversion of leucoanthocyanidin to epicatechin showed constitutive up-regulation when only exposed to the biostimulant. Such priming-related reprogramming of flavonoids in absence of stress was commonly reported in leaves of grapevine treated with seaweed extract^29,30^ or oregano essential oil vapours for instance^39^. Enhanced flavonol accumulation has been previously associated with improved drought tolerance, as demonstrated in *Arabidopsis* lines overexpressing the transcription factors *AtMYB12* and *AtPAP1*^40^, as well as in a maize mutant naturally enriched in flavonols^41^. In the Col-0 *Arabidopsis* background, flavonols modulate root development, acting as negative regulators of root hair formation^42^, lateral root initiation^43^, and root gravitropism through the modulation of auxin transport^44^, most likely via scavenging of reactive oxygen species. However, mutant characterization in other genetic backgrounds revealed a broad spectrum of additional architectural phenotypes in both roots and shoots. In this context, the differential accumulation of flavonoids observed in grapevine roots may have reshaped root system architecture, leading to a moderate reduction in biomass, that remained difficult to quantify in our study (Fig.2b). Crucially, this potential architectural adjustment did not appear to increase susceptibility to stress. On the contrary, it may reflect an adaptive trade-off, where a more compact yet metabolically buffered root system is compensated by enhanced antioxidant capacity, as observed in treated plants, at least during the 2021 season (Fig.5). Unfortunately, the approach used in this study to assess architectural changes did not provide sufficient resolution to clearly conclude on root structural modifications (Fig.2b). For this reason, further targeted studies focusing specifically on root systems will be required to determine whether such architectural changes, potentially driven by phenylpropanoid pathway reprogramming, actively contribute to improved water uptake and, ultimately, enhanced drought adaptation.

### Conclusion

Overall, our findings reveal the effect of the biotimulant AXIOMA Vine on plant physiology within a moderate field-realistic drought scenario, where treated plants maintained higher stomatal conductance and Ψ_PD_ than controls. Under well-watered or extremely stressed conditions, however, its effects were negligible. This context-specific efficiency aligns with the view that biostimulant action is conditional on both environmental constraints and crop physiology, and emphasizes the need for detailed monitoring of plant water status in evaluating their effectiveness. Nevertheless, the study also reveals a potential trade-off, manifested as a moderate reduction of some growth parameter, which all together, appears to confer the substantial benefit of limited susceptibility to moderate water deficit conditions. Finally, analysis of gene expression and metabolism highlight a high representation of ROS and stress hormone-responses such as ABA, during the priming phase with the biostimulant. Given the central role of these particular pathway, considered as critical when plants face multifactorial stress^1^, raises the question of the efficiency of the biostimulant when plants are exposed to other fluctuating weather events, such as flooding or cold stress.

## Material and methods

### Plant based biostimulant

The biostimulant AXIOMA Vigne (MA 1190734) is an assemblage of individual plant extracts, all manufactured by Axioma Biologicals (Brive-la-Gaillarde, France). Extracts were obtained by hydroalcoholic extraction (55,7 % v/v) at room temperature, from aerial parts of plants belonging from families with potential biostimulant properties, namely *Thuya occidentalis* L., *Trigonella foenum-graecum* L., *Solanum dulcamara* L., *Capsicum annum* L., *Matricaria chamomilla* L., *Euphrasia officinalis* L., *Laurus nobilis* L. *Sambucus nigra* L., *Urtica urens* L. and *Equisetum arvense* L^45–51^ and diluted with deionized water to reach 0.12% of dry weight. NMR analysis detected sugars (sucrose, arabinose and fructose), organic acids (acetic acid, malonic acid and formic acid), amino acids (glutamine and tyrosine), phenolic compounds, choline, trimethylamine, an isopropylamine-type compounds as major components of the extract and traces of SO_3_ (0.039 %), Fe (0.039 %), ZN (0.000022 %), Cu (0.0000047 %); Mn (0.000012 %), Co (0.00025 %), B (0.039 %), Mo (0.00035 %), MgO (0.099%). In addition, for solid application a concentrated liquid biostimulant form (973 mL of hydroalcoholic plant extracts and co-formulants per litre) was integrated into *Lithothamnium calcareum* of 2-4 mm size containing 95 % of dry matter to a concentration of 50 mL of concentrated biostimulant per kg of Lithothamne.

### Plant material, growing conditions and treatments

The experiment was conducted in 2021 and 2022 following the same protocol. Grapevine plants (*Vitis vinifera* cv. Cabernet Sauvignon) were propagated in a greenhouse from woodcuttings. After 35 days, rooted cuttings were transferred to 2.8 L pots of sand-soil mixture (Klassman RHP 15 commercial potting mix with70 % fair peat of sphaine, 15 % cold black peat, 15 % pearlite and Danish clay). The plants were watered by subirrigation and fertilised twice a week (nutrient solution N/P/K 20/20/20). Eighty-two-month-old plants with 10–12 leaves were placed outside and were divided into four randomized blocks. During this period, plants were exposed to natural fall and winter conditions and were irrigated continuously with drip irrigation. Biostimulant treatments were performed according to the manufacturer instructions for field applications. At leaf senescence (about two weeks after the plants were transferred outside), the solid treatment was applied to the root system of 40 plants by depositing 1.5 g of the solid treatment on the surface of each pot (equivalent to 10 kg.ha^-1^). Then, during the next growing season, the liquid treatment of an equivalent of 2 L.ha^-1^ was applied with three successive foliar treatments at three phenological stages: at fully expanded leaves (BBCH 11-19), at floral bud emergence (BBCH 57-60), and at the fruit set (BCCH 71) stages. At the same time, 40 other plants received deionized water as a control.

### Application of stress conditions

Two weeks after the last treatment (the 7^th^ of June in 2021 and 14^th^ of June in 2022), the treated and untreated plants were transferred to greenhouses and watered to field capacity by flooding them to their maximum capacity. The pots were then left to drain excess water for half a day. Four environmental conditions were then imposed by combining two temperature regimes and two watering regimes. Plants were equally separated either in a temperature-controlled greenhouse maintained at 27–30 °C or in a non–temperature-controlled greenhouse (40 plants per greenhouse), where the cooling system was turned off, resulting in elevated temperatures. The greenhouse temperature of control and Heat stress plants were continuously recorded. Simultaneously (in each greenhouse system), plants were either irrigated daily (well-watered with a drip irrigation system) or subjected to water deprivation (water deficit). These combinations resulted in four distinct stress conditions: plants grown under controlled temperature and well-watered conditions served as the no-stress control (NS); plants exposed to elevated temperatures while remaining well-watered constituted the heat stress treatment (H); plants maintained at controlled temperature without irrigation were assigned to the drought stress treatment (D); and plants exposed to both elevated temperatures and water deficit represented the combined heat and drought stress treatment (H:D). Plants treated with the biostimulant and control (untreated) were exposed to the four abiotic conditions leading to eight modalities (n=10). Plants were maintained under these conditions for the duration of the stress period prior to physiological measurements and sampling. To prevent an excessive water loss by soil evaporation, pots subjected to water deprivation (D and H:D) were placed into dark plastic bags well fixed around the trunk. To prevent plants under combined stress from reaching the wilting point, pots were weighed every two days from day 10 after stress onset (when Ψ_PD_<-0.6MPa) and until the end of the experiment to allow precise management of water lost through evapotranspiration by supplying the same amount of water lost daily.

### Water status measurements

Grapevine water status parameters were monitored every 5 days using two to four plants per modality (control and treated). Predawn water potential (Ψ_PD_) and midday water potentials (Ψ_l_) were measured in MPa using a pressure chamber (Model 1000, PMS Instrument Co.)^52^. Specifically, Ψ_PD_ measurements were completed before sunrise, whereas Ψ measurements were taken between 12am and 2:00pm. Leaves used at predawn and at midday were fully expanded mature leaves, and at midday exposed to direct solar radiation. After being removed from the branch with a razor blade, leaves were immediately inserted into the DG Meca pressure chamber, with the petiole protruding from the cover. Slow pressure was applied, and the potential value (MPa) was determined using a pressure gauge at the appearance of the water meniscus at the end of the petiole. Stomatal conductance (*gs*, mmol.m^-2^.s^-1^) was measured between 12am and 2:00pm using a LI-600 leaf porometer (LI-COR) operated in auto *gsw* mode. The instrument was calibrated every 30 measurements. Measurements were performed on the abaxial leaf side of the uppermost fully expanded leaf of 10 plant biological replicates. All measurements were performed in mixed sequence to avoid systematic errors between treatments caused by temporary environmental changes.

### Plant physiology monitoring

At the end of the stress period, visual impact of different modalities and stress was assessed at least twice by establishing a visual score from 1 to 5 based on visual cues such as leaf discoloration, dried leaves and apex state (1: No visible stress; 2: Low stress; 3: Moderate stress; 4: High stress; 5: Dead plant, Fig. S1). At the end of the experiment, shoot length was measured. In 2022, the number of internodes and total leaf area was determined by using a reference curve linking the size of the central vein of Cabernet-Sauvignon leaves to the leaf area measured on all leaves with a planimeter (R² = 0.9486) (n=10). After 29 and 35 days of stress applications in 2021 and 2022 respectively, roots were carefully cleaned with deionized water and photographed for root architecture analysis, performed with “ARIA” software^53^. Finally total root fresh weight (FW) was determined. Leaf chlorophyll was measured at the end of the experiment on three leaves per plant using the Dualex Force A clamp.

### Sampling

For sampling, 3^rd^ or 4^th^ leaves from the apex and the roots cleaned with water were harvested between 10 am and 1 pm. The central vein of the leaves was removed with a scalpel, and the remaining lamina immediately frozen and stored and at −80 °C. Five replicates were constituted by pooling the leaves or the roots of two different plants. Leaf and roots pools were ground in powder in liquid nitrogen in a cryogrinder (SPEX FreezerMill 6875).

### Biochemical Characterization and Antioxidant Potential Assay

For biochemical determination, 100 mg aliquots of fresh leaf and root powder were extracted with 10 mL of absolute methanol and agitated for 30 min at 4°C. After centrifugation, the supernatant is removed through speed-vac and pellet was solubilized with 2 mL absolute ethanol. Total flavonoid, were determined by adding 200 µL of methanolic to 60 µL NaNO_2_ 7 %. After 6 min at room temperature 120 µL AlCl_3_ à 10 % is added followed with 120 µL AlCl_3_ à 10 % and 250 mM NaOH. The absorbance was measured at 765 nm and compared to a catechin acid standard curve. Total soluble polyphenols was determined by adding 200 µL of methanolic extract to 1 mL of Folin Ciocalteau reagent (Sigma-Aldrich) diluted 10 times, to which was added 800 µL 7.5 % (w/v) aqueous Na_2_CO^3^. After 30 min at room temperature, the absorbance was measured at 765 nm and compared to a gallic acid standard curve. Antioxidant capacity was assessed using Ferric Reducing Antioxidant Power (FRAP) assay. Briefly, 200 µL of extract were mixed with 1 mL of phosphate buffer 0.2 M (pH 6.6) and 1 mL potassium ferricyanide 1 %. After 20 min at 50 °C, trichloroacetic acid 10 % is added. After centrifugation, 1 mL of supernatant is mixed with 1.2 mL of FeCl_3_ 0.016 %. The absorbance was measured at 700 nm.

### Metabolomic analysis

For metabolomic analysis, soluble metabolites were extracted from 10 mg aliquots of lyophilised leaf and root powders. Two successive extractions were performed at 4 °C with a buffer composed of ethanol (80 %), 10 mM Hepes–KOH (pH 6) with formic acid 0.1 % and the obtained supernatants were pooled together.

Untargeted metabolite profiling of those extracts was carried out using ultra-high-performance liquid chromatography coupled to high-resolution mass spectrometry (UHPLC-HRMS). Specifically, analyses were performed with an Ultimate 3000 UHPLC system (ThermoScientific, Bremen, Germany) connected to an LTQ-Orbitrap Elite mass spectrometer equipped with an electrospray ionization (ESI) source. Data acquisition was conducted in negative ionization modes, following the methodology described^54^. Chromatographic separation was achieved on a C18 Gemini column (2.0 × 150 mm, 3 µm particle size, 110 Å pore size; Phenomenex, Torrance, CA, USA). Full-scan high-resolution MS data were collected at a resolving power of 240,000 (measured at m/z 200). In addition, tandem MS (MS/MS) spectra were obtained in higher-energy collisional dissociation (HCD) mode, using normalized collision energies of 60 %. A pooled Quality Control (QC) sample was generated by combining 20 µL from each individual sample and biological standard. QC injections were performed every fifth run and served to monitor analytical consistency throughout the untargeted metabolomics workflow. These QC data were used to compute the coefficient of variation (CV) for each detected metabolite feature, ensuring that only the most reliable signals were included in subsequent chemometric analyses. Raw LC-MS datasets were processed using MS-DIAL version 4.8^55^ After curation steps—including blank subtraction, signal-to-noise ratio (SN) filtering above 10, and exclusion of features with CVs greater than 30% in QC samples—a total of 3,594 features were retained. Among these, 316 were annotated at Level 2 confidence with both MS1 and MS2 spectral matches, 2,264 features showed MS1-only matches (Level 3 identification), and 1,716 features remained unannotated. For the annotated compounds, InChiKeys were submitted to ClassyFire to assign chemical classification using automated structural ontology.

### RNA extraction and reverse transcription

Three biological replicates of leaf and root per modality were selected randomly for RNA extraction. Fresh leaf powder at 80 mg was added to 12 µL of TCEP and 1.4 mL of extraction buffer preheated to 56 °C (300 mM Tris-HCl, pH 8.0, 25 mM EDTA, 2 M NaCl, 20 g/L cetrimonium bromide (CTAB), 20 g/L polyvinylpolypyrrolidone (PVPP), 500 μL/L tri Spermidine hydrochloride (0.05 %) (≥ 98 % Sigma) and 10 g/L β-mercaptoethanol added extemporaneously). The mixture was shaken and centrifuged at 5600 g for 5 min. The following steps of total RNA extraction were performed using the NucleoMag RNA kit from MACHERY-NAGEL following the manufacturer’s instructions. Total RNA was reverse transcribed using 2 μL-Oligo(dT)_12-18_ (Invitrogen), ribonuclease inhibitor and M-MLV reverse transcriptase (Invitrogen) following the manufacturer’s instructions, cDNAs were stored at −20 °C.

### Gene Expression

Gene expression was assessed using quantitative polymerase chain reaction (qPCR), following the method previously described with the BioStim96 microarray^29^. Building on the existing chip design and incorporating additional genes associated with water stress tolerance in grapevine, a new chip named “VitiSummerGen” was developed. This chip includes a selection of genes from various biological processes: pathogen-related (PR) proteins (n = 12), genes from secondary metabolism including the phenylpropanoid pathway (n = 11), the salicylic acid pathway (n = 2), the mevalonate pathway (n = 1), and the shikimate pathway (n = 4). Additional genes are involved in redox metabolism and homeostasis (n = 10), signaling pathways responsive to heat and drought stress (n = 13), and phytohormonal signaling (auxin, jasmonic acid, abscisic acid; n = 10). Genes related to structural adaptation, such as those involved in cell wall remodeling (n = 8), were also included. Finally, the array encompasses genes linked to physiological functions such as water transport (n = 4), energy metabolism (n = 8), nutrient acquisition (n = 6), and cell/plastid division (n = 5). A full list of genes and associated functions is provided in Table S1.

The specificity of each primer pair was confirmed by checking the size of the amplified product (data not shown) and by the presence of a single melting peak in qPCR analyses. Primer efficiency values ranged from 0.8 to 1.2, allowing relative gene expression to be calculated using the simplified ΔΔCq method (2^−ΔΔCq^) derived from the Pfaffl model. Three housekeeping genes (*VvGAPDH, VvTIP41, VvTHIORLYS8*) were used as internal standards for normalization RT-qPCR was conducted according to the MIQE (minimum information for publication of quantitative real-time PCR experiments) guidelines ^56^, and final relative expression was the ratio from the sample and the average of relative expression control NS from of its respective year and organ.

### Statistical analyses

Statistical analyses were performed using R Software (R Core Team, Vienna, Austria, 2025). Normality and homogeneity of variance were analysed by Kolmogorov–Smirnov and Levene’s tests. The significance of the results was assessed (p-value < 0.05) using independent samples Student’s t-tests, one-way analysis of variance (ANOVA) followed by Duncan’s tests, or two-way ANOVA, as described in figure legends. Density plots were performed with ggExtra packages. For principal component analysis (PCA) the FactoMineR package was used^57^. Data visualization was performed with Excell and ggplot2 package^58^. Determination of Area Under the Curve (AUC) was performed by Trapezoidal Integration with PRACMA package^59^. Venn diagrams were plotted using the online software https://www.interactivenn.net/. Treatment effects on group repartitions for visual stress were tested with Fisher’s test with a 5 % threshold.

## Acknowledgements

The authors gratefully acknowledge the technical assistance provided by Pierre Gastou and Fanny Pinoteau. We also thank Akinao staff for their valuable help in conducting field trials and laboratory analyses. This work received financial support from the French government in the framework of the IdEX Bordeaux University “Investments for the Future” program / GPR Bordeaux Plant Sciences.

## Author contributions

MD, AMB, GC and CELD conceived and designed the experiments. MD and JB performed field and laboratory work. MD and JB carried out transcriptomic and metabolomic analyses. TP, MD and JB performed data analysis and statistical modelling. TP and MD wrote the manuscript with the contribution from GC and CELD. All authors read and approved the final version of the manuscript.

## Data availability statement

All data can be found online in the main text and supporting information materials (Table S2). The metabolic dataset and all metadata will be deposited online after journal acceptance for publication using Dataverse INRAE.

## Supplementary information

**Fig. S1:** Contribution of different root architectural parameters to PC1 and PC2 in Principal component analysis (PCA) of biostimulant treated and control plants under no stress and combined stress conditions in 2021 (a) and 2022 (b).

**Fig. S2:** Illustrative pictures for different stress classification estimated at the end of the stress phase.

**Fig. S3:** Hierarchical clustering analysis of genes exclusively affected by Heat, AUC_gs_ and/or a combination of them in leaves and roots.

**Fig. S4:** Hierarchical clustering analysis from the correlation matrix generated with a total of 1364 metabolites in leaves and 956 metabolites in roots.

**Fig. S7:** Putatively annotated phytohormone related-metabolites under biostimulant (Blue) or control conditions (Grey) under No Stress and Heat:Drought conditions in roots and leaves.

**Supplementary Table S1. :** Names, pathway categories and references of the 96 genes used for the VitiSummerGen microarray chip.

**Supplementary Table S2. :** Data file

