## Supplementary Figures for "A plant-based biostimulant modulates grapevine susceptibility within a realistic water stress window through priming and phenylpropanoid pathway regulation"

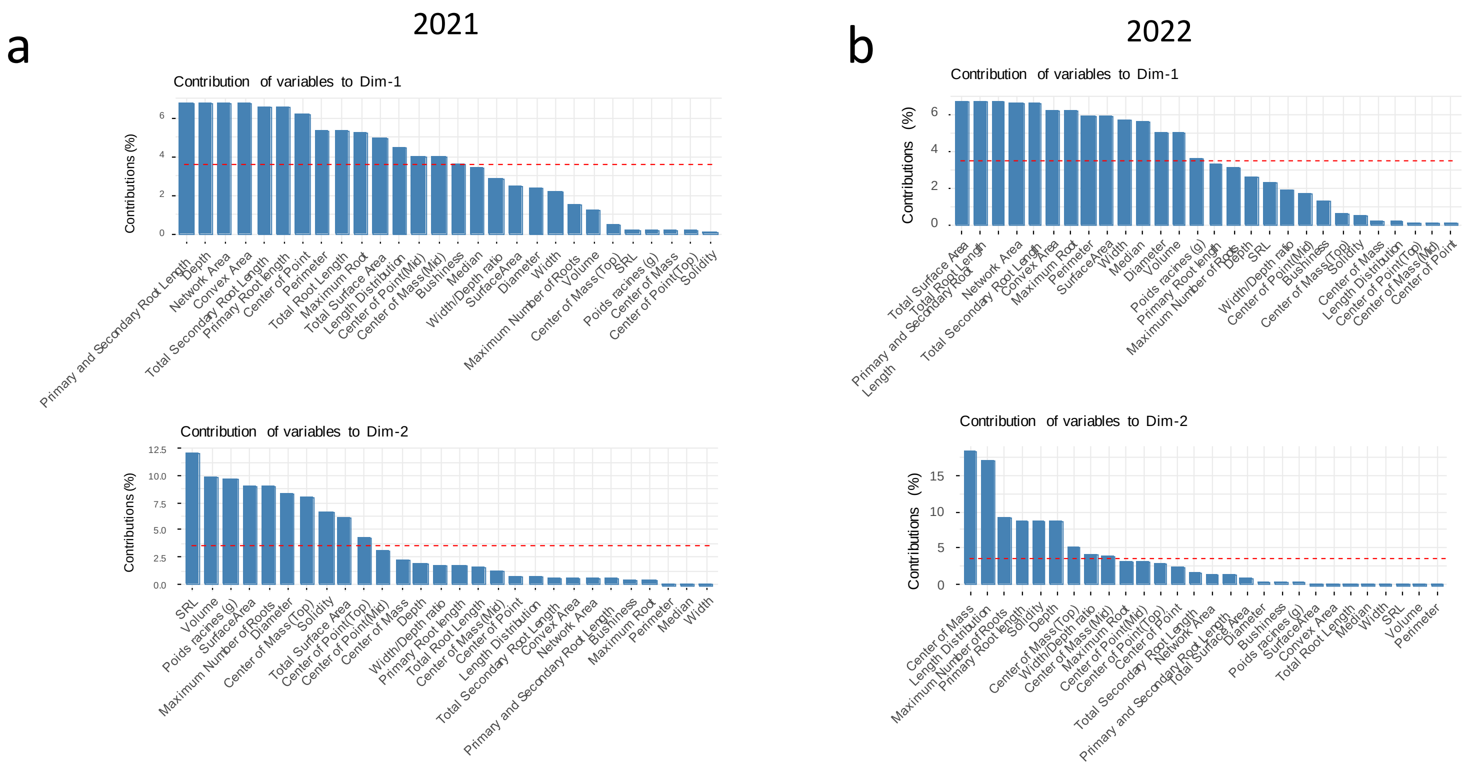


**Fig. S1:** Contribution of different root architectural parameters to PC1 and PC2 in Principal component analysis (PCA) of biostimulant treated and control plants under no stress and combined stress conditions in 2021 (a) and 2022 (b).

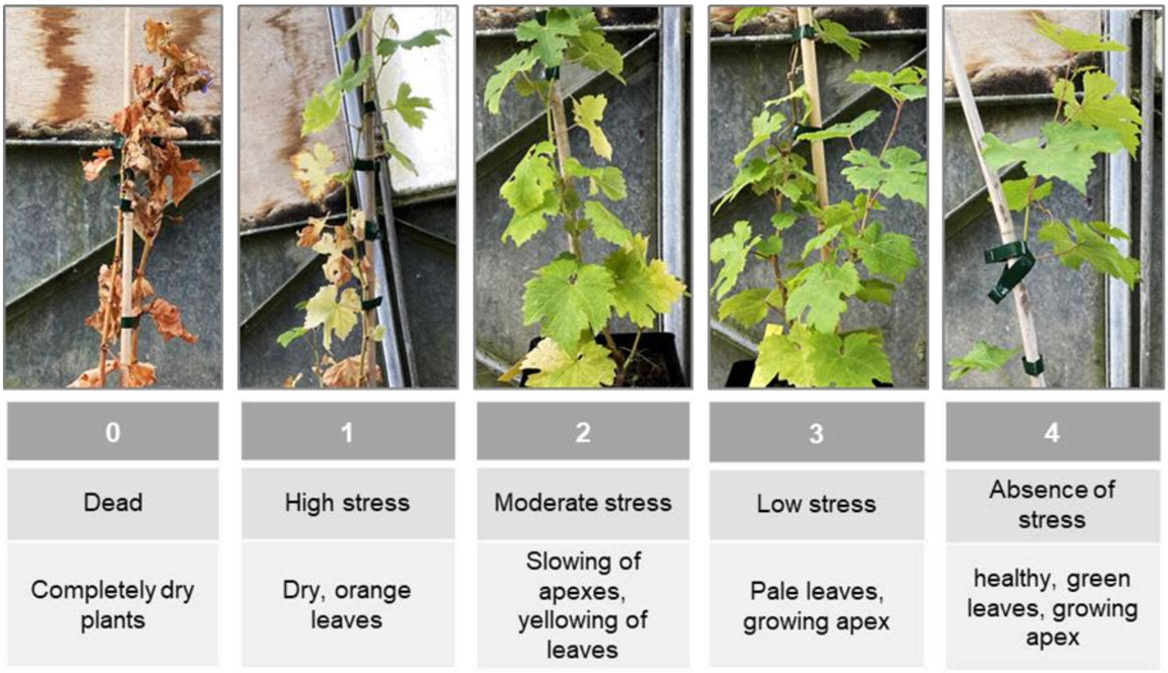


**Fig. S2:** Illustrative pictures for different stress classification estimated at the end of the stress phase (0: Dead plant ; 1: High stress; 2: Moderate stress ; 3: Low stress; 4: No visible stress. Criteria are based on visual cues such as leaf discoloration, dried leaves and apex state.

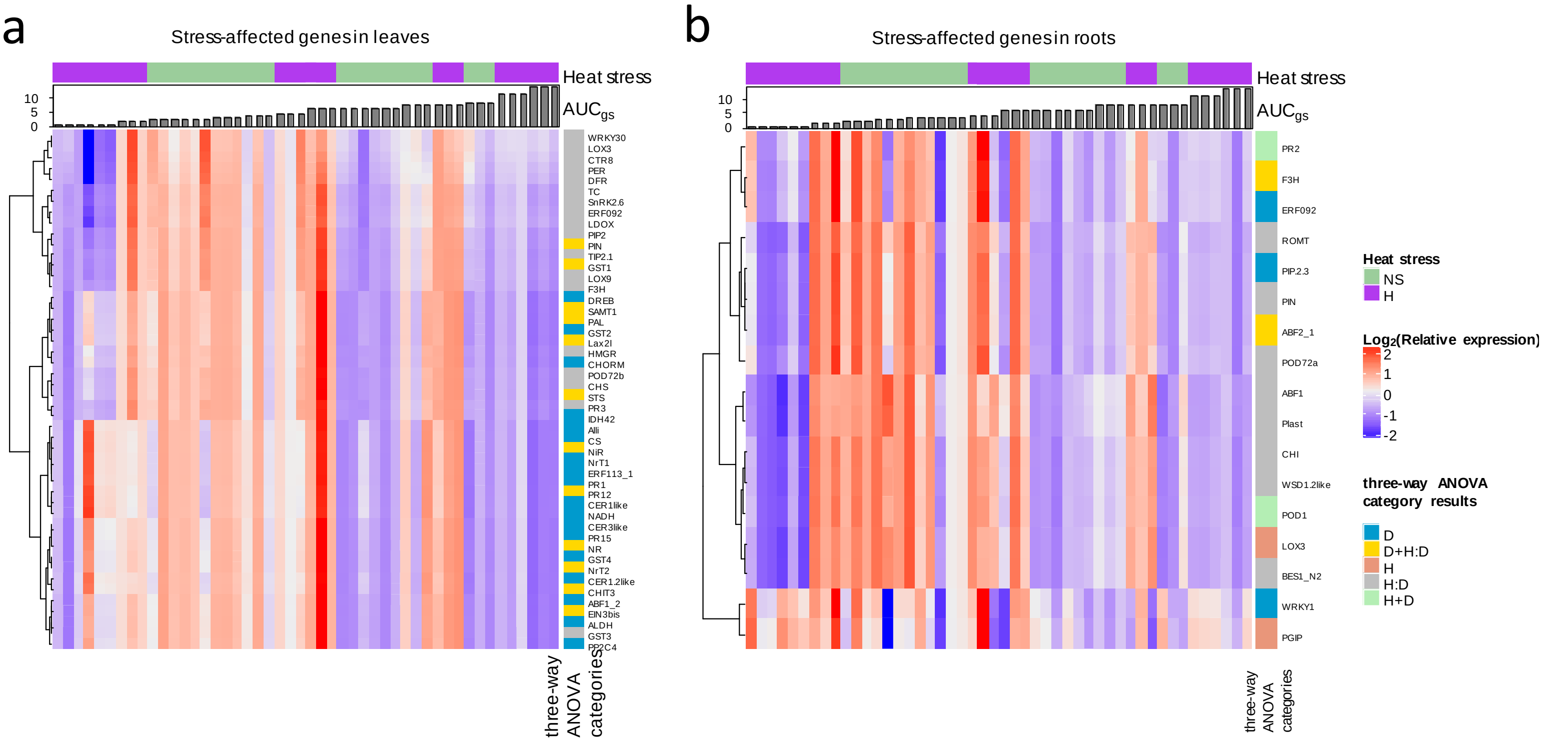


**Fig. S3:** Hierarchical clustering analysis of genes exclusively affected by Heat, AUC$_g$ and/or a combination of them in leaves (a) and roots (b), according to three-way ANOVA analysis (Drought:Heat:Treatment; $p<0.05$). Samples were ordered according to their AUC$_{gs}$ values shown on the top of the heatmaps. Upper colored bar indicates heat stress modality (Green: No heat stress; Purple: Heat stress). Color scale (red to white through blue) indicates the centered and reduced value of log$_2$(Relative expression) of genes of interest. Color bar next to gene names indicates the three-way ANOVA category results, namely: D:Drought; H:Heat; H:D: Interaction of Heat and Drought; H+D Additive effect of Heat and Drought; D+H:D: Additive effect of Drought and the interaction of Heat and Drought;

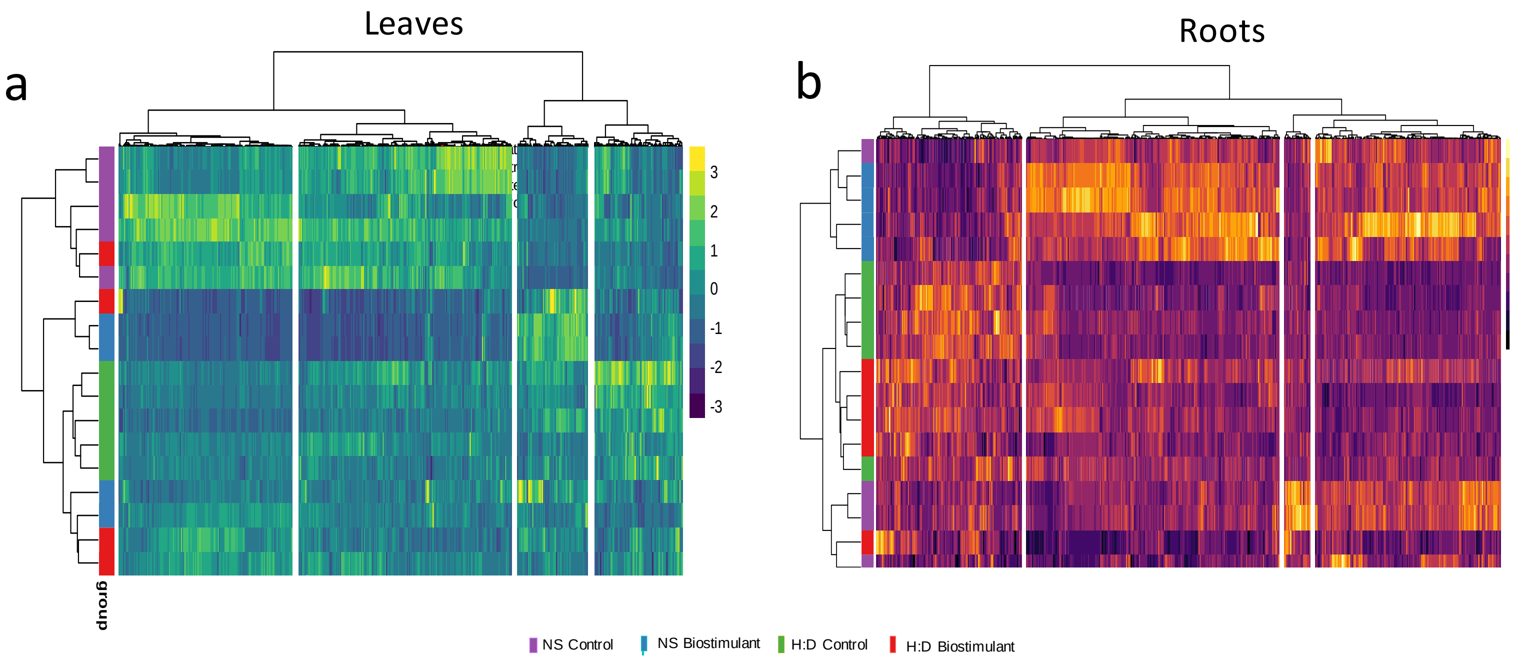


**Fig. S4:** Hierarchical clustering analysis from the correlation matrix generated with a total of 1364 metabolites in leaves (a) and 956 metabolites in roots (b), to evaluate the individual and combined effects of treatment and H:D stress according to three-way ANOVA, (Drought:Heat:Treatment; $p<0.05$). Color scales (Yellow to blue for leaves and Yellow to red for root) indicates the centered and reduced value of $log_2$(Relative expression) of different features. Colored scale at the left of the heatmaps indicates the modalities of the clustered samples (Purple: NS control; Blue: NS Biostimulant; Green: H:D Control; H:D Biostimulant)

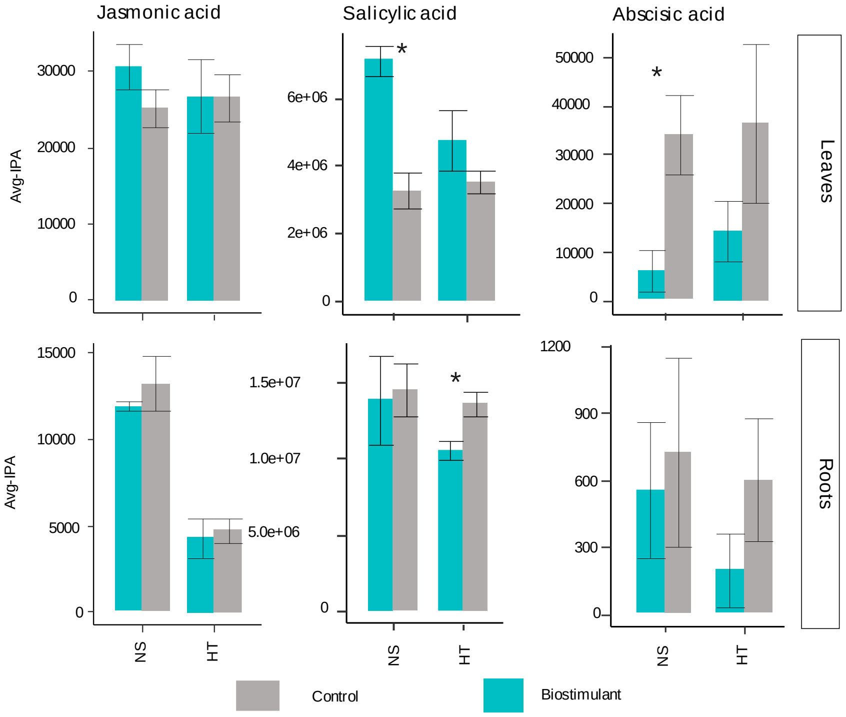


**Fig. S5:** Putatively annotated phytohormone related-metabolites under biostimulant (Blue) or control conditions (Grey) under NS and H:D conditions in roots and leaves. Value represent mean ± se (n=4-5). Significant differences between control and treated plants are highlighted with asterisks (* ≤ 0.05).
