## Supplementary Tables S1 for "A plant-based biostimulant modulates grapevine susceptibility within a realistic water stress window through priming and phenylpropanoid pathway regulation"

TableS1 : Names, pathway categories and references of the 96 genes used for the VitiSummerGen microarray chip

| Metabolic pathway | *Name* | Gene Function | Accession Number | Reference | PCR Efficacy |
|---|---|---|---|---|---|
| Signalling | *VvBES1_N2* | Protein brassinazole-resistant 1 | XM_002283316.3 | Bao Gu et al., 2020 (1) | 0.96 |
| | *VvDREB* | DREB | XM_002276117.5 | Lixia Hou et al.2020 (2) | 1.16 |
| | *VvEIN3* | Ein3-binding F box protein 1 | XM_02285213.1 | Dufour et al (2016) (3) | 0.98 |
| | *VvERF092* | Function of ethylene responsive factor gene family | XM_002282145.5 | Xiaoming Sun et al., 2019 (4) | 0.81 |
| | *VvERF113_1* | AP2/ERF domain-containing protein | VIT_01s0150g00120 | Xiaoming Sun et al., 2019 (4) | 0.88 |
| | *VvNAC 17* | Transcription factors | | Lingye Su et al., 2020 (5) | 0.96 |
| | *VvPP2C4* | probable protein phosphatase 2C 68 | XM_002280432.4 | Yu-Ting Wang et al., 2019 (6) | 1 |
| | *VvWRKY1* | Transcription factors | AY585679.1 | Dufour et al (2016) (3) | 1 |
| | *VvWRKY13_2* | Transcription factors | XM_002278988.4 | Lixia Hou et al.2020 (2) | 0.9 |
| | *VvWRKY30* | Transcription factors | NC_012022.3 | Dan Zhu et al., 2018 (7) | 1.01 |
| | *VvMAPKK1* | Vitis vinifera mitogen-activated protein kinase kinase 1 mRNA | MT154080.1 | Gang Wang et al, 2020 (8) | 0.8 |
| | *VvSnRK 2.6* | Serine/threonine-protein kinase SAPK10 | XM_003631568.3 | Yu-Ting Wang et al., 2019 (6) | 1 |
| Hormons signaling | *VvABA2* | Xanthoxin dehydrogenase | XM_002265688.3 | Bodin et al (2020 (9) | 0.83 |
| | *VvABF1* | GRIP55 protéine bZIP | VIT_18s0001g10450 | Yu-Ting Wang et al., 2019 (6) | 1 |
| | *VvABF1_2* | GRIP55 protéine bZIP | XM_010665762.2 | Yu-Ting Wang et al., 2019 (6) | 0.83 |
| | *VvABF2_1* | Abscisic acid-insensitive 5-like protein 5 | VIT_03s0063g00310 | Yu-Ting Wang et al., 2019 (6) | 0.83 |
| | *VvABF2_2* | Abscisic acid-insensitive 5-like protein 5 | XM_010649445.2 | Yu-Ting Wang et al., 2019 (6) | 1 |
| | *VvALDH* | Aldehyde dehydrogenase family 3 member F1 | XM_002273322 | Bodin et al (2020 (9) | 1.01 |
| | *VvLax2l* | Auxin transporter-like protein 2like | XM_002277381.3 | Bodin et al (2020 (9) | 0.91 |
| | *VvLOX3* | Lipoxygénase | XM_002284499.2 | Dufour et al (2016) (3) | 1.09 |
| | *VvLOX9* | Lipoxygénase | AY159556 | Dufour et al (2016) (3) | 0.92 |
| Homeostasis | *VvPOD72 (a)* | *Vitis vinifera* peroxidase 72 | XM_002275273.4 | Huilin Xiao et al., 2020 (10) | 0.84 |
| | *VvPOD72 (b)* | *Vitis vinifera* peroxidase 72 | XM_002275273.4 | Huilin Xiao et al., 2020 (10) | 0.7 |
| | *VvPODN1* | *Vitis vinifera* peroxidase N1 | XM_003633318.3 | Huilin Xiao et al., 2020 (10) | 0.85 |
| | *VvPOD1* | *Vitis vinifera* cationic peroxidase 1 | XM_002285687.2 | Huilin Xiao et al., 2020 (10) | 1.15 |
| | *VvPOD1bis* | *Vitis vinifera* cationic peroxidase 1 | XM_003634432.3 | Huilin Xiao et al., 2020 (10) | 1.3 |
| Redox status | *VvGST1* | GST (glutathione S-transferase - GST) | AY156048.1 | Dufour et al (2016) (3) | 0.99 |
| | *VvGST2* | GST (glutathione S-transferase - GST) | AY156049 | Dufour et al (2016) (3) | 0.84 |
| | *VvGST3* | GST (glutathione S-transferase - GST) | XM_002283178 | Dufour et al (2016) (3) | 1.06 |
| | *VvGST4* | GST (glutathione S-transferase - GST) | XM_002271673 | Dufour et al (2016) (3) | 1 |

| | | | | | |
|---|---|---|---|---|---|
| | *VvRafS1* | Raffinose synthase | VIT_05s0077g00840 | Hongrui Wang et al. 2020 (11) | 0.81 |
| | *VvFTSH2* | ATP-dependent zinc metalloprotease FTSH 6 | XM_002283357.4 | Valerie Farai Masocha et al., 2020 (12) | 0.83754 |
| | *VvFTSH6* | ATP-dependent zinc metalloprotease FTSH 2 | XM_019223684.1 | Valerie Farai Masocha et al., 2020 (12) | 0.94283 |
| | *VvGSS1 x1* | Granule-bound starch synthase 1, chloroplastic/amyloplastic | XM_019225517.1 | Valerie Farai Masocha et al., 2020 (12) | 1.02 |
| | *VvMDH* | Malate deshydrogenase | XM_002278676.3 | Bodin et al (2020 (9) | 1.28 |
| | *VvNADH* | | | Dufour et al (2016) (3) | 0.95 |
| | *VvCS* | Citrate synthase | XM_002271415.2 | Bodin et al (2020 (9) | 0.97 |
| | *VvCP 29.1* | Chlorophyll a-b binding protein CP29.1, chloroplastic | XM_002279798.3 | Valerie Farai Masocha et al., 2020 (12) | 0.96 |
| | *VvIDH42* | Isocitrate Deshydrogénase | XM_002270581.3 | Bodin et al (2020 (9) | 1.08 |
| | *VvPDV1* | Plastid division protein | XM_002276562.4 | Bodin et al (2020 (9) | 0.87 |
| | *VvATP At* | AAA-ATPase | XM_002268977.4 | Valerie Farai Masocha et al., 2020 (12) | 0.85 |
| | *VvPlast* | Plastidic aldolase protein | TC52601 | Bodin et al (2020 (9) | 0.96 |
| | *VvTC* | Transketolase, chloroplastic | XM_002280724.4 | Valerie Farai Masocha et al., 2020 (12) | 1.1 |
| | *VvNiR* | Nitrite Réductase | NM_001281265.1 | Bodin et al (2020 (9) | 1.01 |
| | *VvNR* | Nitrate Réductase | NM_001281120.1 | Bodin et al (2020 (9) | 1.02 |
| | *VvNrT1* | Nitrite/nitrate transporteur | KF649633.1 | Bodin et al (2020 (9) | 1.18 |
| | *VvNrT2* | Nitrite/nitrate transporteur | XM_002277091.2 | Bodin et al (2020 (9) | 1.2 |
| | *VvSUC 27* | Vitis vinifera sucrose transporter-like | NM_001281141.3 | Cai Yumeng et al., 2020 (13 - 14) | 0.89 |
| | *VvCTR8* | Copper transporter | HQ108192 | Bodin et al (2020 (9) | 0.87 |
| Plant growth | *VvCYC* | G2/mitotic-specific cyclin | XM_002283116.3 | Bodin et al (2020 (9) | 0.92 |
| | *VvPAL* | Phénylalanine ammonialyase | XM_002268220.1 | Dufour et al (2016) (3) | 1.1 |
| | *VvSTS* | Stilbene synthase (resvératrol synthase) | X76892.1 | Dufour et al (2016) (3) | 0.95 |
| | *VvCHI* | Chalcone Isomérase | *X75963* | Dufour et al (2016) (3) | 1.04 |
| | *VvCHI2* | Chalcone Isomérase 2 | XM_002280122 | Dufour et al (2016) (3) | 1.12 |
| | *VvCHS* | Chalcone Synthase | X75969.1 | Dufour et al (2016) (3) | 1.01 |
| | *VvCHS2* | Chalcone Synthase 2 | XM_002276885.1 | Dufour et al (2016) (3) | 1.04 |
| | *VvLDOX* | Anthocyanidine synthase | X75966 | Dufour et al (2016) (3) | 0.84 |
| | *VvDFR* | DFR | XM_002281822.1 | Dufour et al (2016) (3) | 1.05 |
| | *VvF3H* | Flavanone-3-hydroxylase | *X75965.1* | Dufour et al (2016) (3) | 0.87 |
| | *VvCHORM* | Chorismate mutase | *FJ604854* | Dufour et al (2016) (3) | 1.05 |
| | *VvCHORM2* | Chorismate mutase 02 | XM_002284083.1 | Dufour et al (2016) (3) | 1.11 |
| | *VvCHORS* | Chorismate Synthase | *FJ604855* | Dufour et al (2016) (3) | 1 |
| | *VvCHORS2* | Chorismate synthase 1 | XM_002282263 | Dufour et al (2016) (3) | 1.11 |
| | *VvROMT* | Resvératrol O-methyl-transferases | FM178870 | Dufour et al (2016) (3) | 0.96 |
| | *VvSAMT1* | SA Methyl Transferase | XM_002262982.1 | Dufour et al (2016) (3) | 1.16 |
| | *VvHMGR* | 3-hydroxy-3-methylglutaryl Coenzyme A reductase | XM_002275791.1 | Dufour et al (2016) (3) | 0.93 |
| Secondary metabolism (defenses) | *VvPR1* | PR1 Unknown function | AJ536326 | Dufour et al (2016) (3) | 1 |

| | | | | | |
|---|---|---|---|---|---|
| | *VvPR10* | PR protein - class 10 (PR10) | *AJ291705* | Dufour et al (2016) (3) | 0.99 |
| | *VvPR12* | Defensin | XM_002281153 | Dufour et al (2016) (3) | 0.88 |
| | *VvPR15* | Germin like Protein | NM_001281199.1 | Dufour et al (2016) (3) | 1.19 |
| | *VvPR2* | Vitis vinifera bêta 1-3 glucanase | XM_002277475 | Dufour et al (2016) (3) | 0.96 |
| | *VvPR3* | Endochitinase (Chitinase IV) [PR3] | U97522.1 | Dufour et al (2016) (3) | 1.16 |
| | *VvPR8* | Vitis vinifera chitinase 1-like | XM_002276329 | Dufour et al (2016) (3) | 0.96 |
| | *VvCHIT3* | Chitinase 3 - PR8 | Z68123 | Dufour et al (2016) (3) | 0.98 |
| | *VvGLU* | Beta-1,3-glucanase (PR2) | AF239617 | Dufour et al (2016) (3) | 1.18 |
| | *VvPOX* | Lignin-forming peroxidase | XM_002285687.1 | Dufour et al (2016) (3) | 1.14 |
| | *VvPGIP* | Polygalacturonase Inhibiting Protein | XM_002263487.1 | Dufour et al (2016) (3) | 0.96 |
| | *VvPIN* | Proteinase inhibitor PR6 | XM_002284418 | Dufour et al (2016) (3) | 1.23 |
| Reinforcement cell wall | *VvWSD1-2 like* | Vitis vinifera O-acyltransferase WSD1 | XM_019225727.1 | Dimopoulos et al., 2020 (15) | 1 |
| | *VvCER1_2 like* | Vitis vinifera protein ECERIFERUM 1 | XM_002263751.4 | Dimopoulos et al., 2020 (15) | 0.88 |
| | *VvCER1_like* | Protéine Vitis vinifera ECERIFERUM 1 | XM_002265153.4 | Dimopoulos et al., 2020 (15) | 0.89 |
| | *VvCER10_like* | CER10 protein (CER10) | VIT_13s0019g01260 | Dimopoulos et al., 2020 (15) | 0.91 |
| | *VvCER3 like* | Vitis vinifera protein ECERIFERUM 3 | XM_002269997.4 | Dimopoulos et al., 2020 (15) | 0.88 |
| | *VvAlli* | Alliinase | XM_002265837.1 | Dufour et al (2016) (3) | 1.04 |
| | *VvPER* | Vitis vinifera hypothetical protein | XM_002274762.1 | Dufour et al (2016) (3) | 0.93 |
| Aquaporins | *VvTIP1-1* | Tonoplast intrinsic protein 1;1 | NM_001280994.1 | Bodin et al (2020 (9) | 1.02 |
| | *VvTIP2-1* | Tonoplast intrinsic protein 2;1 | NM_001280994.1 | Megan C. Shelden et al., 2017 (16) | 1.12 |
| | *VvPIP 2-3* | Plasma membrane Intrinsic Protein 2;3 | NM_001281129.1 | Bodin et al (2020 (9) | 0.99 |
| | *VvPIP2* | Plasma membrane Intrinsic Protein 2 | DQ358107 | Irene Peronne et al., 2012 (17) | 0.93 |


1- Bao Gu, Bo Zhang, Lan Ding, Peiying Li, Li Shen, Jianxia Zhang (2020) Physiological Change and Transcriptome Analysis of Chinese Wild *Vitis amurensis* and *Vitis vinifera* in Response to Cold Stress. **Plant Molecular Biology Reporter**, Volume 38, pages 478–490, (2020)

2- Lixia Hou, Xinxin Fan, Jie Hao, Guangchao Liu, Zhen Zhang, Xin Liu (2020) Negative regulation by transcription factor VvWRKY13 in drought stress of *Vitis vinifera* L. **Plant Physiology and Biochemistry**, Volume 148, Pages 114-121

3- Dufour Marie-Cécile, Magnin Noël, Dumas Bernard, Vergnes Sophie and Corio-Costet Marie-France (2016) High-throughput gene-expression quantification of grapevine defense responses in the field using microfluidic dynamic arrays. **BMC genomics**, 17: DOI 10.1186/s12864-016-3304-z.

4- Xiaoming Sun, Langlang Zhang, Darren C. J. Wong, Yi Wang, Zhenfei Zhu1, Guangzhao Xu, Qingfeng Wang, Shaohua Li, Zhenchang Liang and Haiping Xin (2019) The ethylene response factor VaERF092 from Amur grape regulates the transcription factor VaWRKY33, improving cold tolerance. **The Plant Journal** 99, 988–1002

5- Lingye Su, Linchuan Fang, Zhenfei Zhu, Langlang Zhang, Xiaoming Sun, Yi Wang, Qingfeng Wang, Shaohua Li & Haiping Xin (2020) The transcription factor VaNAC17 from grapevine (*Vitis amurensis*) enhances drought tolerance by modulating jasmonic acid biosynthesis in transgenic Arabidopsis. **Plant Cell Reports**, Volume 39, pages 621–634

6- Yu-Ting Wang, Ze-Ya Chen, Yue Jiang, Bing-Bing Duan, Zhu-Mei Xi (2019) Involvement of ABA and antioxidant system in brassinosteroid-induced water stress tolerance of grapevine (*Vitis vinifera* L.). **Scientia Horticulturae**, Volume 256, 15 October 2019, 108596

7- Dan Zhu, Yongmei Che, Peilian Xiao, Lixia Hou, Yang Guo (2018) Functional analysis of a grape WRKY30 gene in drought resistance. **Plant Cell, Tissue and Organ Culture (PCTOC)**, Volume 132, Pages 449-459

8- Gang Wang, Ying-hai Liang, Ji-yu Zhang & Zong-Ming (Max) Cheng (2020) Cloning, molecular and functional characterization by overexpression in Arabidopsis of MAPKK genes from grapevine (*Vitis vinifera*). **BMC Plant Biology**, 20, Article number: 194

9- Bodin Enora, Bellée, Anthony, Dufour Marie-Cécile, André Olivier, Corio-Costet, Marie-France. (2020) Grapevine Stimulation: A Multidisciplinary Approach to Investigate the Effects of Biostimulants and a Plant Defense Stimulator. **Journal of Agricultural and Food Chemistry** 68 (51): 15085-15096

10- Huilin Xiao, Chaoping Wang, Nadeem Khan, Mengxia Chen, Weihong Fu, Le Guan & Xiangpeng Leng (2020) Genome-wide identification of the class III POD gene family and their expression profiling in grapevine (*Vitis vinifera* L). **BMC Genomics**, Volume 21, Article number: 444

11- Hongrui Wang, Joshua J. Blakeslee, Michelle L. Jones, Laura J. Chapin, Imed E. Dami (2020) Exogenous abscisic acid enhances physiological, metabolic, and transcriptional cold acclimation responses in greenhouse-grown grapevines. **Plant Science**, Volume 293, 110437

12- Valerie Farai Masocha, Qingyun Li, Zhenfei Zhu, Fengmei Chai, Xiaoming Sun, Zemin Wang, Long Yang, Qingfeng Wang, Haiping Xin (2020) Proteomic variation in *Vitis amurensis* and *V.vinifera* buds during cold acclimation. **Scientia Horticulturae**, Volume 263, 109143

13- Yumeng Cai, Jing Yan, Qike Li, Zhefang Deng, Shaoli Liu, Jiang Lu, Yali Zhang (2019) Sucrose transporters of resistant grapevine are involved in stress resistance. **Plant Molecular Biology**, Volume 100, 1-2, Pages 111-132

14- Yumeng Cai, Jing Yan, Wenrui Tu, Zhefang Deng, Wenjie Dong, Han Gao, Jinxu Xu, Nan Zhang, Ling Yin, Qingyong Meng and Yali Zhang (2020) Expression of Sucrose Transporters from *Vitis vinifera* Confer High Yield and Enhances Drought Resistance in Arabidopsis. **International Journal of Molecular Sciences**, Volume 21, 7, page 2624

15- Dimopoulos Nicolas, Tindjau Ricco, Wong Darren, Matzat Till and Haslam Tegan (2020) Drought stress modulates cuticular wax composition of the grape berry. **Journal of Experimental Botany**, Volume 71, 10, Pages 3126-3141

16- Megan C. Shelden, Rebecca Vandeleur, Brent N. Kaiser and Stephen D. Tyerman (2017) A Comparison of Petiole Hydraulics and Aquaporin Expression in an Anisohydric and Isohydric Cultivar of Grapevine in Response to Water-Stress Induced Cavitation. **Frontiers in Plant Science**, Volume 8, 1893

17- Irene Perrone, Giorgio Gambino, Walter Chitarra, Marco Vitali, Chiara Pagliarani, Nadia Riccomagno, Raffaella Balestrini, Ralf Kaldenhoff, Norbert Uehlein, Ivana Gribaudo, Andrea Schubert, Claudio Lovisolo (2012) The Grapevine Root-Specific Aquaporin VvPIP2;4N Controls Root Hydraulic Conductance and Leaf Gas Exchange under Well-Watered Conditions But Not under Water Stress. **Plant Physiology**, Volume 160, Issue 2, Pages 965–977
